# Seasonal niche differentiation between evolutionary closely related marine bacteria

**DOI:** 10.1101/2020.12.17.423265

**Authors:** Adrià Auladell, Albert Barberán, Ramiro Logares, Esther Garcés, Josep M. Gasol, Isabel Ferrera

## Abstract

Bacteria are highly dynamic in marine environments, where they play key biogeochemical roles. Here, we tested how similar the niche of closely related marine bacteria is and what are the environmental parameters modulating their ecological responses in a coastal oligotrophic time series. We further explored how conserved the niche is at broader taxonomic levels. We found that, for certain genera, niche similarity decreases as nucleotide divergence increases between closely related amplicon sequence variants, a pattern compatible with selection of similar taxa through habitat filtering. Additionally, we observed evidence of niche partitioning within various genera shown by the distinct seasonal patterns of closely related taxa. At broader levels, we did not observe coherent seasonal trends at the class level, with the order and family ranks conditioned to the patterns that exist at the genus level. This study explores the coexistence of niche overlap and niche partitioning in a coastal marine environment.

## Introduction

Marine microbial communities are highly dynamic and variable over time, particularly in temperate coastal environments. Community structure changes on a daily, monthly and annual scale due to bottom-up factors such as resource availability (including inorganic nutrients and dissolved organic carbon), top-down biotic interactions and physical properties such as temperature or day length (Fuhrman et al., 2015). The combination of all these factors defines the ecological niches in which microbes grow and reproduce depending on the metabolic potential of each taxa. Given that microbes are key players in the functioning of the biosphere, understanding how taxa adapt to these conditions and respond to environmental changes is crucial (Falkowski, 2012).

Our capability to study the composition of microbial communities has improved drastically during the last decades due to the DNA sequencing revolution. High throughput sequencing of the 16S rRNA gene has facilitated the assessment of microbial diversity in samples collected during global expeditions or from long-term monitoring stations (Buttigieg et al., 2018; Logares et al., 2020; Sunagawa et al., 2015). Notably, microbial observatories have provided time series of several consecutive years, in a crucial effort to extract robust patterns of these microbial assemblages and their dynamics. Early studies using fingerprinting methods and clone libraries had already pointed out an effect of seasonality on the whole bacterioplankton community structure (Alonso-Sáez et al., 2007; Chow et al., 2013; Cram et al., 2015); these methods however only allowed to recover the most abundant taxa. The use of massive sequencing substantially increased the throughput, generating massive amounts of sequence data that were grouped into Operational Taxonomic Units (OTUs) through sequence clustering (usually at an arbitrary cutoff often established between 97 to 99%) helping to reduce data volume, which in turn compensated for possible sequencing errors (Callahan et al., 2017). Alongside sequencing technology improvements, new bioinformatic algorithms have increased the level of resolution at which we can analyze sequence data by allowing to work with amplicon sequence variants (ASVs), differentiating up to one nucleotide difference (Callahan et al., 2016). The delineation of sequence variants has shown how an OTU can contain variants with different ecological behaviors likely representing different species or ecotypes (Callahan et al. 2016). For example, Eren et al. (2013) showed how the method could differentiate between two SAR11 ecotypes with only two nucleotide differences of the 16S rRNA gene that displayed anti-correlated seasonal patterns. Likewise, Chafee M. et al. (2018) showed recurrent switching of ecotypes at single nucleotide resolution during spring and summer phytoplankton blooms driven by temperature and substrate changes. In addition, studies focused on the potential association between photosynthetic picoeukaryotes and bacteria have shown how the use of ASVs has improved the association signal by identifying stronger correlations among them (Lambert et al., 2018; Needham et al., 2018).

Hutchinson proposed that an ‘n-dimensional hypervolume’ could define the niche of a species: a set of conditions under which an organism can survive and reproduce, including both biotic and abiotic factors (Hutchinson, 1957). Bacteria have adapted to the different conditions present in the marine environment through processes of selection and speciation. If two taxa occupy identical niches, a taxon should eventually outcompete the other; yet in practice, many closely related taxa coexist (Cohan, 2017). The niche would be determined both by the homogeneous selection of traits to survive in a specific environment –e.g. resistance to high salinity, an example of habitat filtering– and the heterogeneous selection for other traits to reduce competition –i.e. niche partitioning– that would facilitate coexistence. In closely related taxa, their distribution can inform on whether two taxa display a similar realized niche –the abiotic conditions together with the interaction of biotic factors such as competition– or if ecotype differentiation occurred through niche partitioning. In this sense, time series of marine microbial observatories are useful for identifying taxa with similar realized niches through co-occurrence analyses with repeated sampling over time (Friedman & Alm, 2012). Additionally, and while niches are commonly considered as features of species, we can extend the definition of the Hutchinsonian niche to broader taxonomical groups and evaluate the importance of the shared traits within each group and their responses to the environment (Tromas et al., 2018). Such an analysis at different taxonomical levels concur with studies that discuss the importance of the ‘phylogenetic scale’ (Ladau & Eloe-Fadrosh, 2019; Martiny et al., 2015) at which ecology operates. For marine bacteria, it is unclear how niche similarity and the seasonal trends are distributed at wider taxonomic levels such as family, order or class. Yet, the methodology required to address these questions is nowadays available.

Here, we used time series data from a coastal marine observatory in the NW Mediterranean to describe the long-term seasonal trends in bacterial community structure. First, we focused on determining niche similarity between ASVs within genera and later extended the comparison to broader taxonomic levels to answer (1) how many ASVs are seasonal and what is the temporal distribution of the relevant taxonomic groups, (2) how similar the niche between closely related ASVs within different marine genera is and what are the environmental parameters modulating their distinct ecological responses, and (3) how conserved the realized niche is as we move from genus to broader taxonomic levels (i.e., family, order and class).

## Results

### Environmental, ecological and taxonomic context

Surface water temperature at Blanes Bay varied seasonally, with minimal mean values in February (12.6°C) and maximal values in August (24.5°C, Supplementary Figure 1). Inorganic nutrients were higher during autumn and winter while Chlorophyll *a* reached the highest values (ca. 1 mg·m^−3^) during the winter-spring period. A detailed description of the seasonality at Blanes Bay, including these and other environmental parameters, can be found in Gasol et al. (2016).

In the 11 years of monthly data, we detected a total of 6,825 ASVs. The ASV distribution was compared both by occurrence (narrow, intermediate and broad) and abundance (abundant, rare; see Material and Methods). Most of the them (91%) displayed a narrow distribution, occurring in less than 10% of the samples (Figure 1A, Table 1). Only 26 ASVs displayed a broad distribution (≥75% occurrence), 3 of them always belonging to the rare fraction (i.e. <1%). Taxonomically, among the broad ASVs, 19 belonged to the Alphaproteobacteria, mostly to the orders Pelagibacterales (13 ASVs) and HIMB59 (4 ASVs; former SAR11 clade V; See Supplementary Table 1 for the ASV taxonomic information and Supplementary Table 2 for the correspondence between GTDB and SILVA nomenclature). The 506 ASVs with intermediate occurrence (<75% and >10% occurrence) belonged taxonomically to 20 different classes. The dominant classes were the Alphaproteobacteria and Gammaproteobacteria (163 and 133 ASVs respectively) followed by Bacteroidia (106 ASVs), mostly the Flavobacteriales order (91 ASVs; Figure 1A). The ASVs with a narrow distribution displayed a similar taxonomic composition. We also evaluated the ASVs that were rare but occasionally became abundant (Conditionally Rare Taxa, CRT, see Material and Methods) and found a total of 81 ASVs that met this criterion. Gammaproteobacteria (48 ASVs) and Alphaproteobacteria (13) were the most common CRTs, while the rest belonged to the Verrucomicrobiae and Bacteroidia classes (Figure 1B).

**Figure 1:**
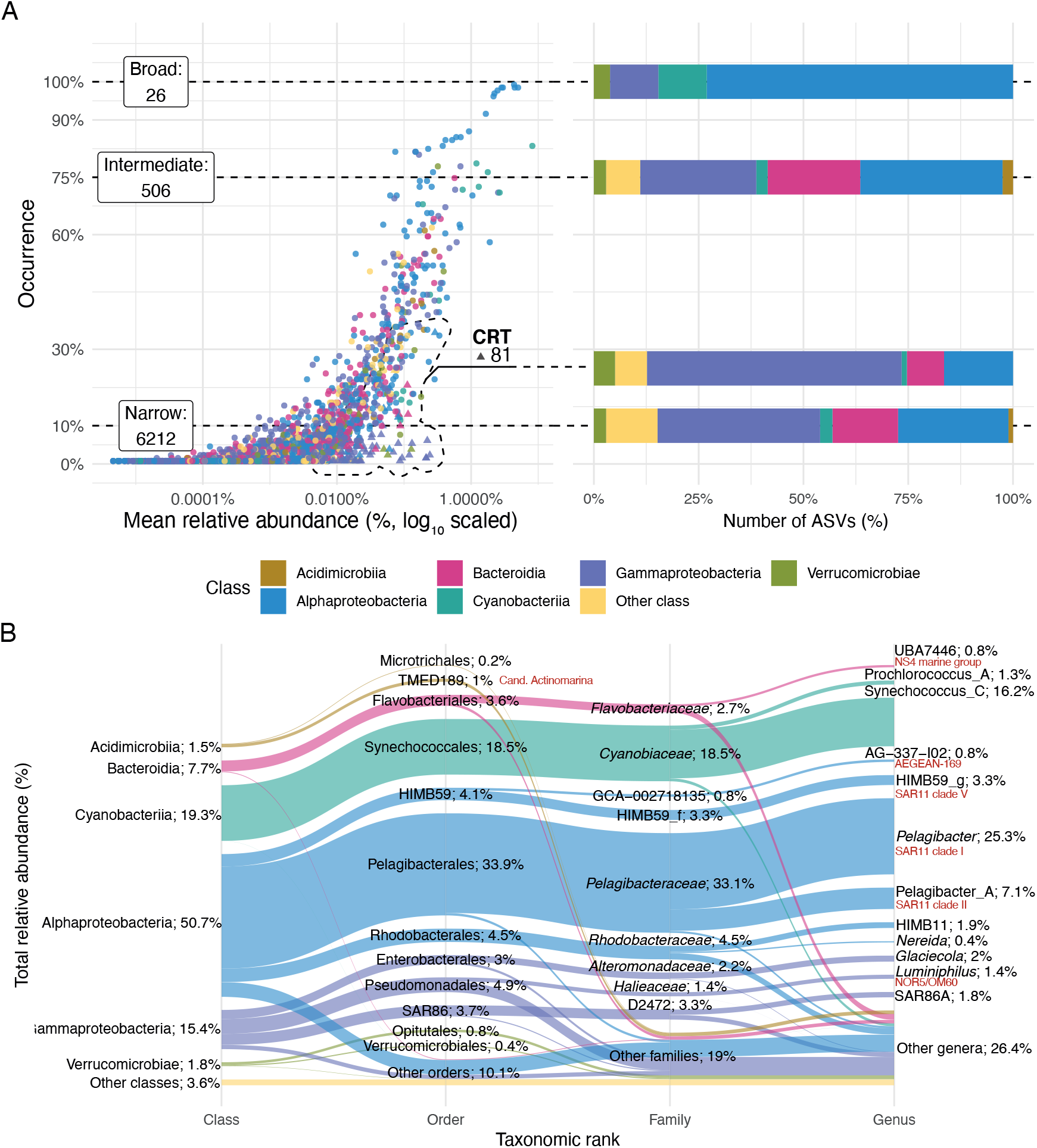
A) Distribution of the different ecological ASVs types (broad, narrow or intermediate, and conditionally rare taxa, CRT). The X axis indicates the occurrence (% of samples) and the Y axis corresponds to the mean relative abundance (%) over the time series. Dotted lines delimitate the distributions (in the label the numbers of ASVs of each type are displayed) and connect to a box indicating the number of ASVs for each distribution and a bar plot colored by taxonomy at the class rank. B) Alluvial plot showing the total relative abundance distribution of Blanes Bay taxa across different taxonomic ranks (class, order, family and genus). The height of the sections displays the relative abundance (indicated in the text; the total is 100%). The SILVA nomenclature is displayed in red next to the corresponding GTDB database nomenclature, used in the text in those cases in which there is no similarity in the names.

**Table 1:**
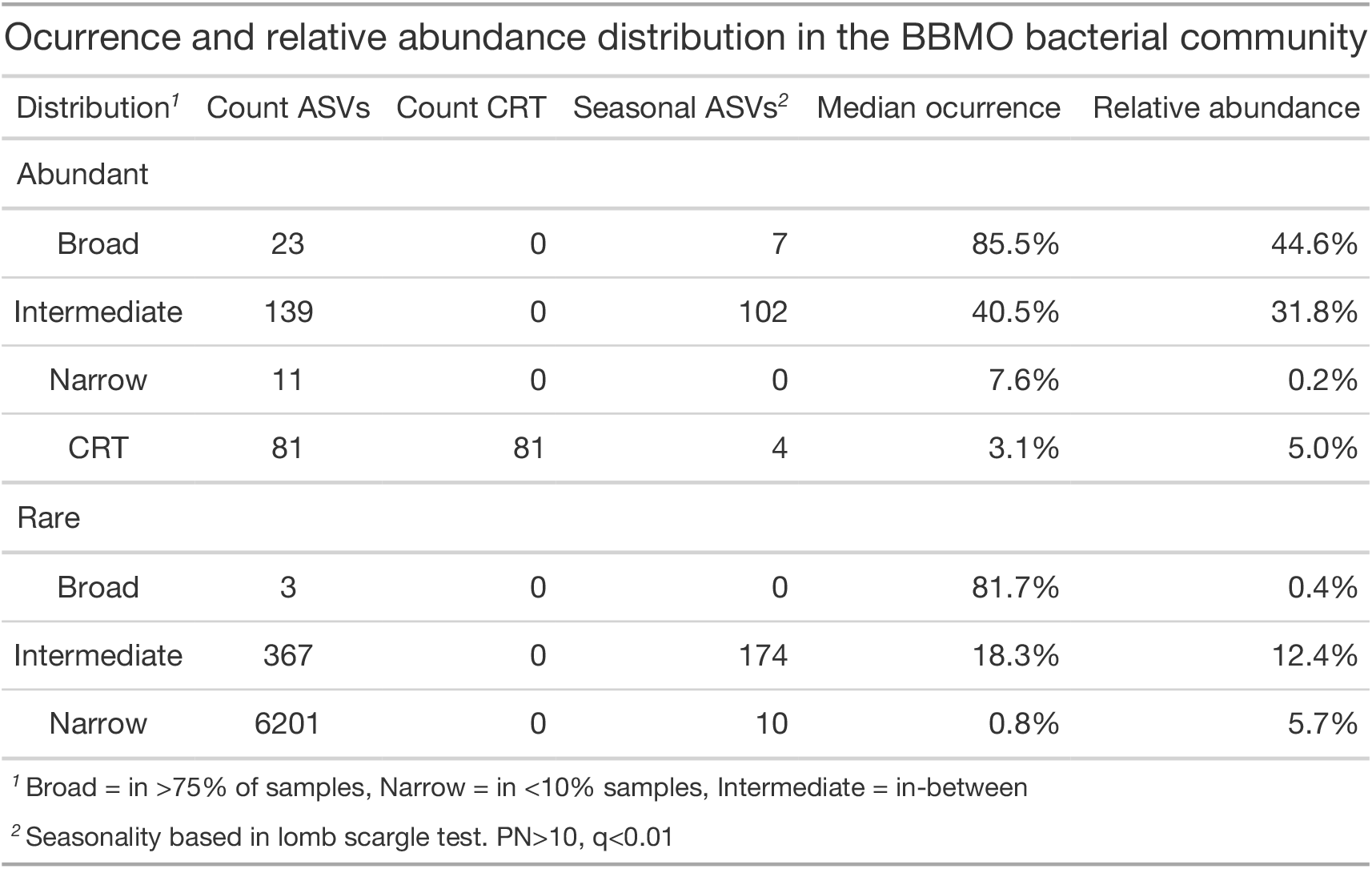
Occurrence and median relative abundance for the ASVs in the Blanes Bay Microbial Observatory dataset. Distribution specifies the occurrence distribution: broad (≥75% samples), narrow (<10% samples) and intermediate. The results are distributed between abundant (≥1% in at least one sample) and rare ASVs. Count ASVs stands for the number of ASVs; Count CRT, the number of Conditionally Rare Taxa; seasonal ASVs, the count of seasonal ASVs (based in lomb scargle test, *q* ≤ *0.01*, PN ≥ 10); median occurrence, the % of samples in which the ASVs appears; total relative abundance of the group.

Spring and summer displayed less alpha diversity than autumn and winter (α richness estimates = 197 vs 334 ASVs respectively, *p* < 0.01; Supplementary Figure 2). When checked at the month level, with January as intercept, we observed a significant decrease in richness starting in April (232 ASVs, *p* = 0.015) to regain higher values in October (316 ASVs, *p* = 0.87). Regarding beta diversity (i.e. community similarity), the seasons with the maximal dissimilarity were summer and winter (β Bray Curtis estimate = 0.48, standard error = 0.036), being autumn and spring the ones with the lowest difference (β estimate = 0.21, standard error = 0.047; Supplementary Figure 3), with similar ranges for all the other comparisons.

### ASV seasonality

A total of 297 ASVs displayed high seasonality (lomb scargle test *q* ≤ 0.01, PN ≥ 10) with different ranges of occurrence and season maxima. These seasonal ASVs represented on average 47% of the relative abundance, partitioned in 13 % of the abundance from ASVs exhibiting broad distribution, 34% of intermediate occurrence and 0.1% of narrow presence. In our study, significant peak normalized power values – a statistic that measures how strong is the recurrence– ranged between 10 and 43.1. The highest values corresponded to ASVs with distributions that recurrently presented a peak in one specific season. Examples of this pattern are ASV122, ASV55 and ASV131, belonging to the Acidimicrobiia, Bacteroidia and Alphaproteobacteria classes respectively (Supplementary Figure 4). These ASVs appeared mostly during winter and fall and were absent from spring and summer. Within the seasonal ASVs, we differentiated 3 significantly different clusters (Supplementary Figure 5). The first group, composed of 23 ASVs, includes most of the broadly distributed ASVs that peaked during summer and autumn. Taxonomically, this cluster was composed for the most part of ASVs from *Cyanobiaceae* and *Flavobacteriaceae*. The second cluster, with 30 ASVs, includes those ASVs that peaked during winter and spring, mainly *Pelagibacteraceae* ASVs. Interestingly, this cluster includes an understudied group, *Marinisoma,* that displayed a winter trend in all its seasonal ASVs (5 out of 9 ASVs). Finally, the last cluster was composed of 244 ASVs that presented a less clear seasonal trend likely due to their lower occurrence and relative abundance along the decade, with no dominance of any particular taxonomic group.

Out of the 297 seasonal ASVs, we identified 131 ASVs that could be clustered into 42 OTUs at 99% identity. We found examples of different behaviors within various genera. For example, *Pelagibacter* was represented by 20 different OTUs; 3 of them were composed of only seasonal ASVs, 6 OTUs contained both seasonal and non-seasonal ASVs, and 9 OTUs consisted only of non-seasonal ASVs. Similar trends were observed for other genera such as SAR86A and *Luminiphilus*. On the other hand, we found that niche partitioning was not common, with only 20% of the OTUs displaying seasonal ASVs with clear partition between seasons. In total, 8 ASVs displayed such behavior; that is, seasonal ASVs within 5 nucleotide differences, displaying relative abundances with opposed seasonal trends or with different temporal patterns (see some examples in Figure 2). Most of these patterns could be classified into either an almost complete temporal separation (e.g. ASV48 vs ASV30 within OTU30, affiliated to Puniceispirillales; Figure 2) or restriction of the “temporal” niche (one of the ASVs is only present in a specific month or season although the other is also present; e.g. ASV285 vs ASV337 within OTU243, affiliated to HIMB59). In fact, seven out of 8 ASVs displayed the latter pattern of seasonal restriction.

**Figure 2:**
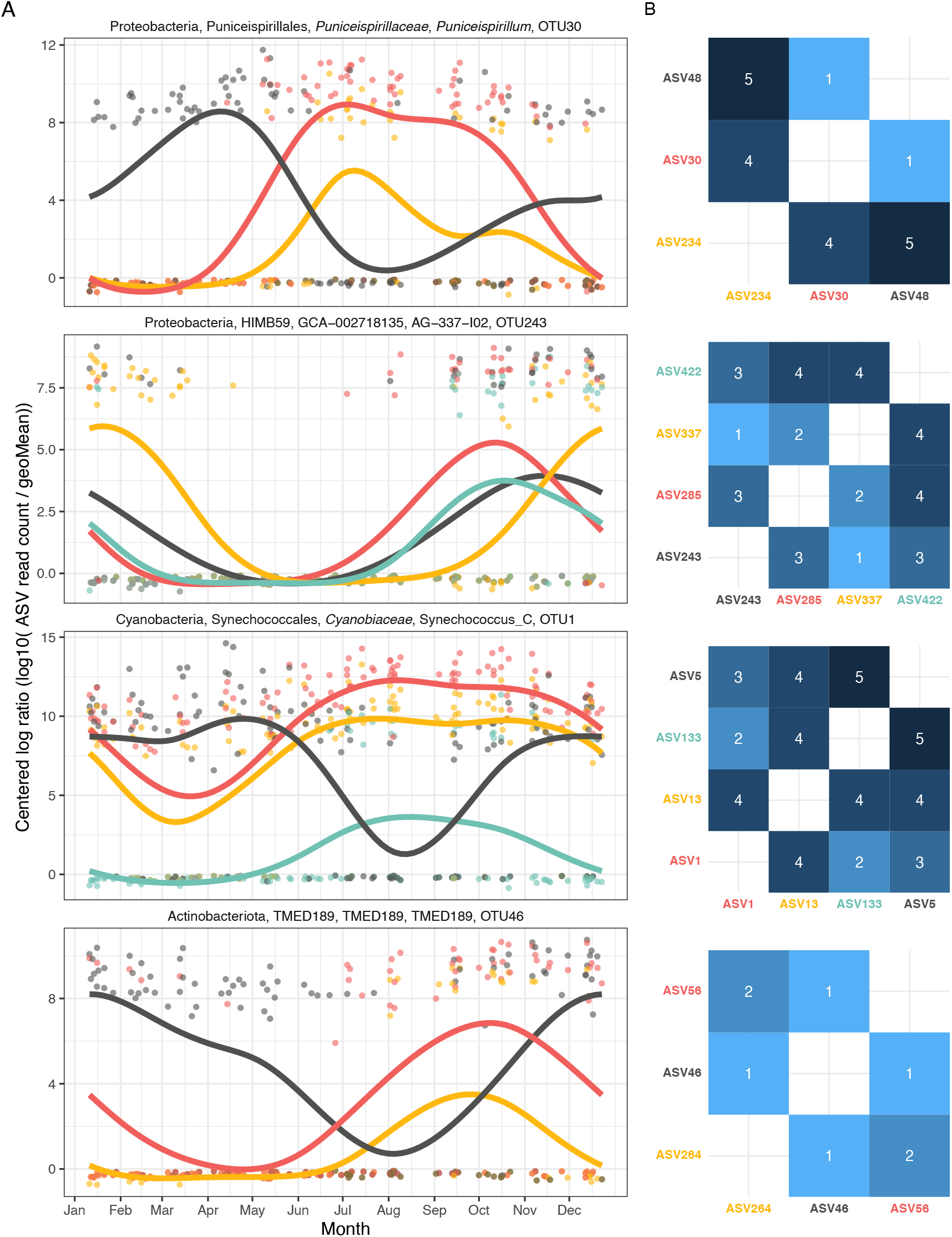
A) Examples of niche partitioning among closely related ASVs within the same OTU (99% clustering). The X axis presents the month and the Y axis presents the centered logarithm ratio abundance. A generalized additive model smooth is adjusted to the data points. B) Heatmaps presenting the nucleotide divergence between each of the ASVs (number of mismatches after alignment). Five nucleotide divergence equals to a median sequence identity of 98.8%.

### Variability of niche preference within genera

Here, we define the ecological niche of a taxon as the set of conditions (biotic and abiotic factors) that fluctuate recurrently in this marine temperate coastal environment and that allow the growth of the organism or its persistence. Taxa display niche preferences, and hypothetically, closely related taxa should have similar niches (Cohan, 2017). A similar ecological niche in two taxa would be represented by the shared environmental conditions that vary over time. In the case that niche overlap exists, cooccurrence and covariance would point to niche similarity, and exclusion situations would indicate the opposite condition, i.e. niche partitioning. Our proxy to test for niche overlap among closely related taxa is the Rho measurement (proportional change between two taxa, see Material and Methods), that can be expressed as a function of the nucleotide divergence (number of nucleotide substitutions between two sequences after an alignment). A decrease in Rho with nucleotide distance means that the taxa decrease their covariance, and therefore behave less similarly as they become more phylogenetically distinct.

Out of the 13 genera evaluated, we found that *Pelagibacter* (Alphaproteobacteria, SAR11 clade I), Pelagibacter_A (Alphaproteobacteria, SAR11 clade II) and SAR86A (Gammaproteobacteria, a subclade of SAR86) displayed a significant decrease in Rho proportionality with increasing nucleotide divergence (Figure 3; See Supplementary Table 3 for the regression statistics). The linear tendency between Rho and the nucleotide distance explained on average about 13% of the trend in Rho. The distributions within each genus were highly variable. The *Pelagibacter* genus displayed the highest number of ASVs (60) and the variation in the Rho score was likewise the highest, between 0.996 and 0.3. The Pelagibacter_A genus presented less ASVs (26) than *Pelagibacter* but a similar Rho distribution. The SAR86A had a smaller amount of variation along the nucleotide change, with a maximum Rho of 0.85. Besides the 3 abovementioned genera, *Luminiphilus* (OM60/NOR5 clade) also displayed a negative tendency but the relationship was not statistically significant. The *Synechococcus* genus displayed similarly high proportionality values at low and high nucleotide distances, not showing a decreasing trend. Merging all the non-significant groups, the values did also not present a significant tendency (data not shown), suggesting that the decrease is specific of some groups.

**Figure 3:**
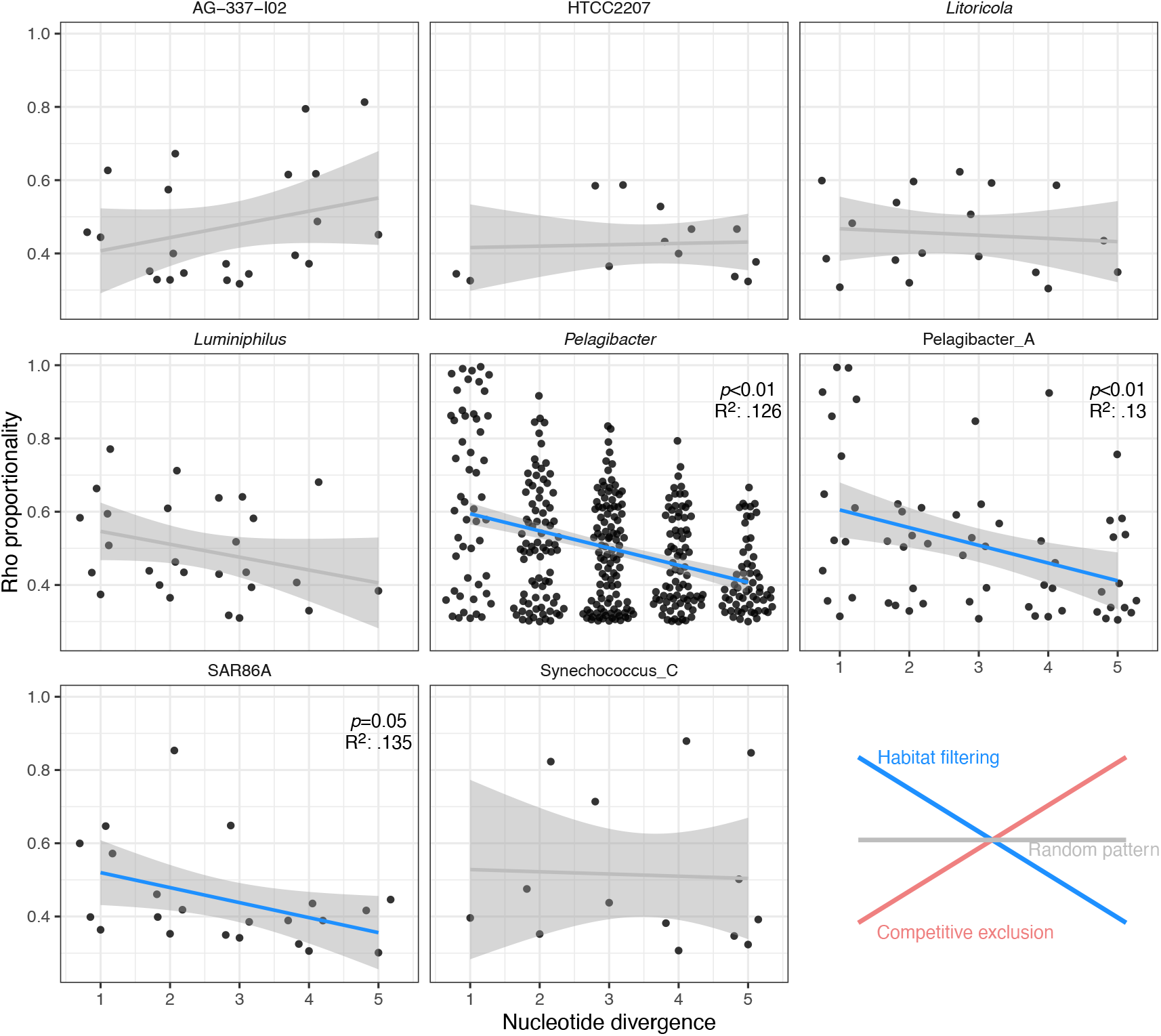
Relationship between the proportionality of change (Rho, Y axis) and the nucleotide divergence (mismatches after alignment, X axis). Only genera with more than 3 ASVs at less than 5 nucleotide divergence were evaluated. Lines represent the linear relationship between the two variables. The blue color indicates statistical significance. The *p* value and the R^2^ are displayed for the significant regressions. Bottom right: a graphical visualization of the different potential ecological patterns. See Supplementary Table 2 for the correspondence between GTDB and SILVA nomenclature.

### Environmental drivers of the observed niche differences within genera

Given the identified differences in the temporal niche (i.e. the time of the year when the organism develops) among closely related ASVs, we further evaluated how different environmental parameters influenced the observed distributions. For each ASV-parameter pair we generated a model and estimated coefficient indicating how the ASV responded (increase or decrease in abundance, Figure 4, Supplementary Figure 6). A total of 245 response models between ASV abundances and environmental parameters out of the 603 possible were significant (FDR ≤ 0.05). About two-thirds of the models were polynomial with the rest being linear. Temperature, nitrite and nitrate concentrations were the parameters appearing most often in significant models, followed by the abundance of photosynthetic and heterotrophic nanoflagellates. The different bacterial genera displayed variability in the responses to the various parameters. *Pelagibacter*, AG-337-I02 (AEGEAN-169 marine group), D2472 (SAR86) and *Luminiphilus* genera had ASVs that responded cohesively, i.e. that displayed the same response sign to a given environmental variability for all their ASVs (Supplementary Figure 6). Most of these bacterial genera showed a negative relative abundance response to temperature and a positive relationshipwith the concentration of inorganic nitrogen compounds. The exception to this trend was *Luminiphilus*, with the opposite coefficient sign for all parameters tested. HIMB59 (former SAR11 clade V), Pelagibacter_A, SAR86A and *Synechococcus* showed differences within each genus, pointing to the existence of distinct ecotypes (Figure 4A). Temperature was a main factor determining these ecotype differences. Within SAR86A, two contrasting patterns could be observed; ASV34 and ASV63 (nucleotide divergence of 1; Supplementary Figure 7) presented a positive relationship to temperature and a negative one to nitrate and chlorophyll *a* concentration, while ASV562, ASV270, ASV65 and ASV157 presented the opposite responses (these ASVs had nucleotide distances ranging from 1 to 9; Figure 4A). In the case of *Synechococcus*, a similar trend was observed (ASV5 and ASV12 *vs.* ASV1 and ASV13, Figure 4) but the nucleotide distances do not hint to a possible explanation based on phylogenetic distance, a result coincident with that of the previous section in which no decrease in niche similarity for this group was observed (Figure 3). Pelagibacter_A also presented two ecotype-specific responses, with ASV6 and ASV10 (1 nucleotide divergence) responding similarly, in contrast to the other significant ASVs within the genus (Figure 4). Finally, the different ASVs belonging to HIMB59 (former SAR11 clade V) presented multiple responses, pointing to a differentiated ecotype distribution.

**Figure 4:**
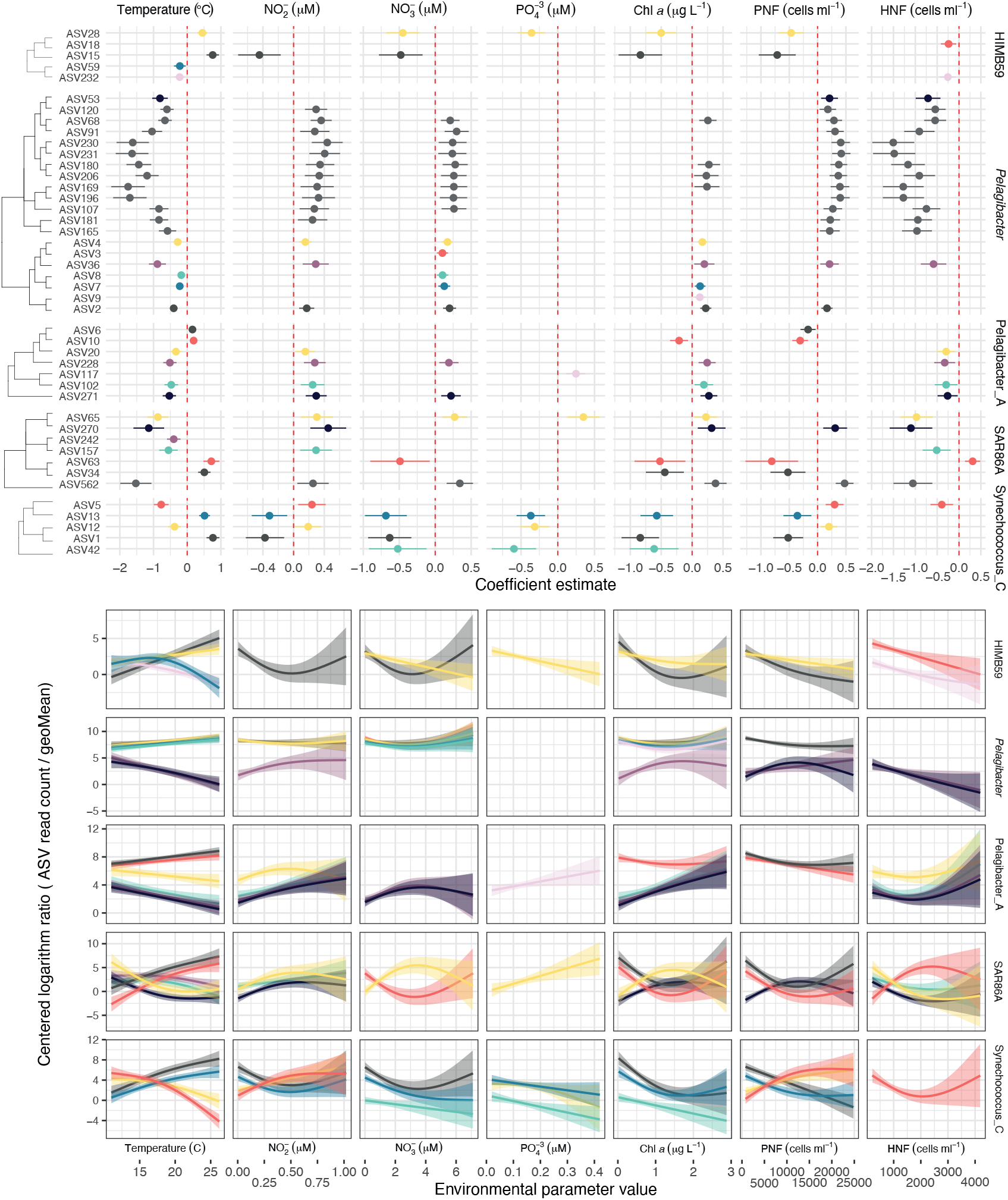
A) Significant *corncob* models between ASVs from HIMB59, *Pelagibacter*, Pelagibacter_A, SAR86 and *Synechococcus* genera (rows) and various environmental parameters (columns). The coefficient estimate indicates positive or negative responses to the parameter and is shown with a 95% confidence interval. The color corresponds to the different ASV within a genus (only the top 8 more abundant ASVs are colored, the other ASVs are shown in grey). ASVs are ordered through a hierarchical clustering based on nucleotide divergence. B) Generalized additive model fits between the ASV centered logarithm ratio abundances and the parameter value distribution for the significant ASVs in the upper plot. Panels and ASV colors shown as in the upper plot. PNF: Phototrophic nanoflagelates; HNF: Heterotrophic nanoflagelates.

### Seasonality at broad taxonomical levels

Having delineated how the ASVs behave seasonally and what are the drivers of the differences within each genus, we tested whether synchronized responses at higher taxonomic levels exist. Theoretically, cohesiveness should decrease from the genus to higher taxonomic ranks. We randomly aggregated 80% of the ASVs at the genus, family, order and class levels to test how this seasonal statistic was distributed. We only considered Alphaproteobacteria, Gammaproteobacteria and Bacteroidia since these were the classes with enough representation down to the genus rank (only levels with >10 ASVs were considered). When we analyzed the general distribution across ranks, we found that the class rank was mostly non-seasonal (98.9% PN values, *p* < 0.01, PN < 10; Figure 5). Both the order and family ranks displayed a similar distribution with ~50% of the results being seasonal, while this value increased up to ~60% at the genus rank. These distributions were different for each class, with Alphaproteobacteria presenting a clear bimodality while Gammaproteobacteria presented values evenly distributed across the PN statistic (Figure 5). By checking each level separately, the bulk Alphaproteobacteria class distribution (Supplementary Figure 8, PN mean = 5.3) could be linked directly to that of the Pelagibacterales order, since this was the most abundant group (Supplementary Figure 8B) and appeared as non-seasonal (PN mean = 5.7, Supplementary Figure 8A). Observing the other prevalent orders (Rhodobacterales, Puniceispirillales –SAR116 clade– and HIMB59), the seasonality statistic was quite robust when randomly removing different ASVs (Supplementary Figure 8). Puniceispirillales for example appeared mostly during summer. This observation was different for the Gammaproteobacteria orders (Supplementary Figure 9A), with SAR86 and Pseudomonadales orders close to the seasonality threshold resulting in half of the randomizations as non-seasonal. Moreover, for the Pseudomonadales order, we observed that it was composed of various families, each with different seasonality (Supplementary Figure 9B). The Bacteroidia class only showed seasonality at the genus level for UBA7446, a new unknown genus within the family *Flavobacteriaceae* (Supplementary Figure 10). Thus, we observed that the distributions at the order level were diametrically different, with Alphaproteobacteria including orders that were seasonal, Gammaproteobacteria orders presenting a peak in the limit of seasonality and all orders of Bacteroidia presenting a non-seasonal trend. Nevertheless, in most groups the family and genus ranks presented similar seasonal trends to those displayed by the order they belonged to.

**Figure 5:**
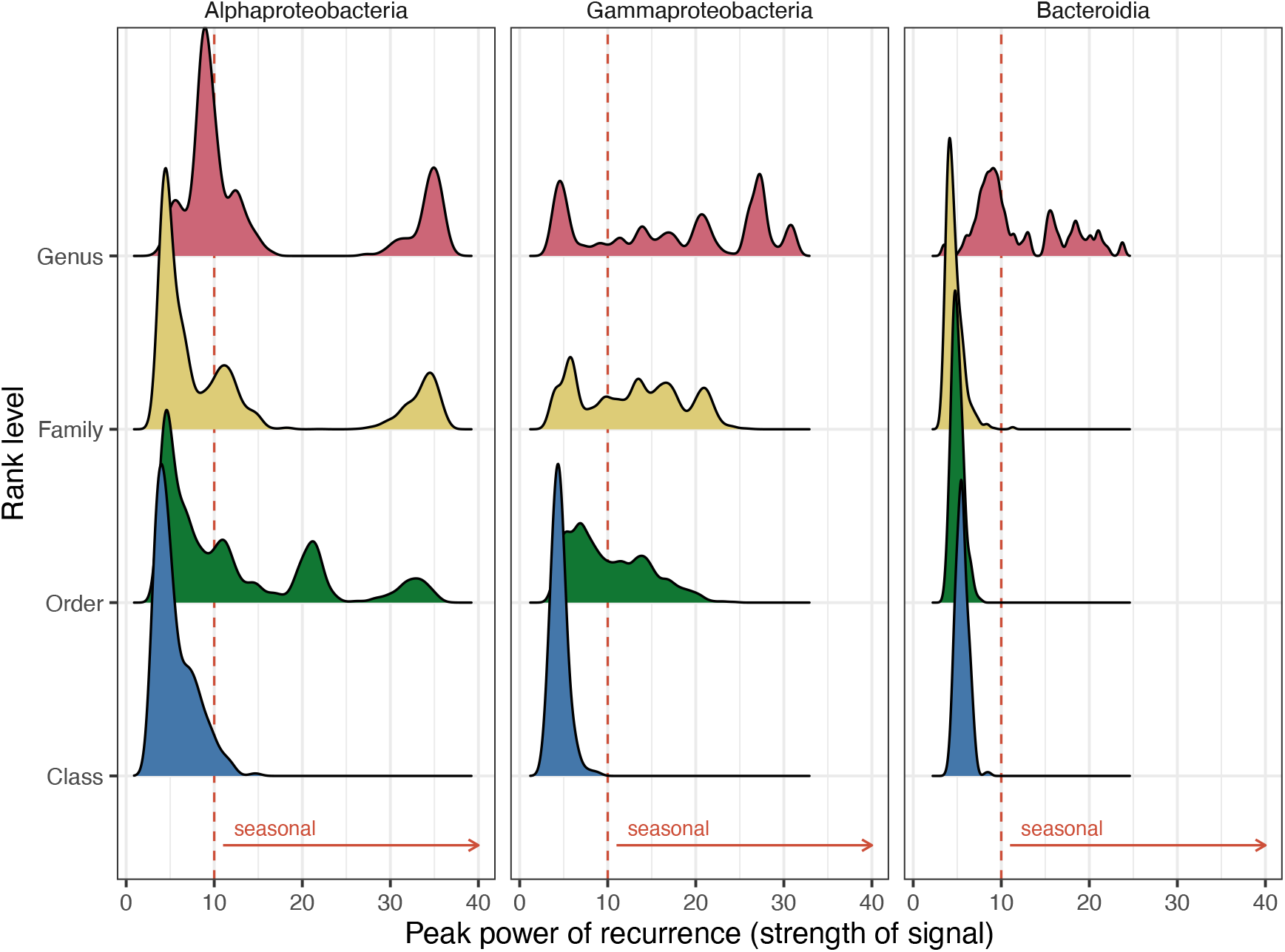
Density distribution of the peak normalized power statistic (as proxy for seasonality) for each rank level in the Alphaproteobacteria, Gammaproteobacteria and Bacteroidia classes. The red line indicates the used threshold for seasonality (*q* ≤ 0.01 and PN ≥ 10).

## Discussion

We explored how the bacterial community is structured seasonally at fine taxonomical levels and whether the structure is maintained at broader levels through long-term sampling and amplicon sequencing in a temperate marine coastal environment. Specifically, we wanted to understand how closely related ASVs respond to the environmental conditions that appear recurrently in the site. Overall, our results show that around half of the total community relative abundance shows seasonality at the ASV level. Within genus, we show how niche similarity decreases with increasing nucleotide divergence for at least 3 genera, while other trends were observed in other groups. We then checked how various environmental parameters define the niche for the components of various genera. Finally, we analyzed how the patterns of seasonality aggregate at the broader taxonomic ranks, showing that, in our dataset, the class levels were non-seasonal and that the other ranks tested (i.e. order and family) present a variety of trends.

Before further considerations, a methodological limitation must be discussed. The use of amplicon marker gene has its limitations for the delineation of biological units (VanInsberghe et al., 2020). The use of hypervariable regions of the 16S rRNA gene –in this case the V3-V4 regions– entails problems regarding the level of taxonomic resolution that can be determined. In fact, VanInsberghe et al. (2020) showed that for *Vibrio* sp. only 7 out of 14 species were distinguishable with 100% full length 16S rRNA gene sequence, implying that a shorter region would be even less informative. The power of the 16S rRNA gene to resolve closely related taxa changes for different bacterial clades, but in general, various studies have shown that the variable regions have a poor resolution for full species delineation (Johnson et al., 2019; VanInsberghe et al., 2020). Nevertheless, despite the abovementioned limitations, amplicon marker gene sequencing still represents the fastest and most comprehensive approach for studying ecological patterns through identifying robust trends in large datasets. To stay on the conservative side in our interpretations, we set the genus level as the one for which we can assign patterns with some certainty.

### Contrasting environmental conditions throughout the year

The environmental parameters displayed a clear seasonal pattern, with the highest rates of change between the summer and winter periods, and the bacterial community mirrored these changes as observed in alpha and beta diversities. The patterns of alpha and beta diversity were studied before at our study site but in much shorter surveys (1-2 years; Alonso-Sáez et al. 2007; Mestre et al. 2017). The analysis of eleven years of data unveiled that the highest differences in community structure occur between summer and winter, and the highest variability is found in spring and winter, which could be related to the idiosyncratic phytoplankton blooms that occur during these periods, with differing intensity over the decade (Nunes et al. 2018; see also PNF in Supplementary Figure 1).

In the nearby long-term microbial station SOLA (Banyuls-sur-Mer), a seven-year seasonal study was performed comparing the bacterial, eukaryotic and archaeal community through ASV delineation (Lambert et al., 2018). The number of ASVs in the bacterial community was similar to that observed in this study (6825 ASVs in this study vs 6242 at SOLA) and a similar community composition was observed, for e.g. both Pelagibacteraceae and Synechococcales dominated the communities (Figure 1, Lambert et al. 2018), with *Pelagibacter*, Pelagibacter_A and Synechococcus_C being the most prevalent organisms. However, some differences were detected; a relevant group in our study was the HIMB59 order, initially considered part of the SAR11 clade V (Martijn et al., 2018; Viklund et al., 2013), which was remarkably absent in the SOLA study (Lambert et al., 2018; Salter et al., 2015). This result could be either the reflect of a different taxonomic assignation or related to primer biases. In fact, this group has been assigned a variety of names and phylogenetic positions; the MAGs within the HIMB59 order were identical at the 16S rRNA level with what was previously described as the AEGEAN-169 marine group, which is present in surface and deep waters in a variety of coastal sites (Alonso-Sáez et al., 2007; Cram et al., 2015), while in the SILVA classification AEGEAN-169 appears within the Rhodospirillales order. Martijn et al. (2018), however, concluded that the HIMB59 and other relevant MAGs conform a separate clade neither within the Pelagibacterales nor the Rhodospirillales, in agreement with the Genome Taxonomy Database assignation used here. Previous studies may have pooled HIMB59 into groups other than SAR11 clade V, hiding its presence. Another difference was the presence of SAR11 clade IV, not detected in our study but present in SOLA. Other relevant groups present in this study at Blanes Bay were *Candidatus* Actinomarina, a group within class Acidimicrobiia with small cells (Ghai et al., 2013), *Glaciecola* and HIMB11 (*Roseobacter* clade), all of them representing ≥1% of the total relative abundance.

### Half of the total community is seasonal

Determining seasonality is not trivial, as it implies to take a binary decision for a trait that is likely continuous in a gradient rather than into two states. In our analysis, we found a total of 297 seasonal ASVs (34% of the evaluated ASVs, which made up a total of 47% of the sequences). A lower value was observed by Giner et al. (2019) in a 10-year study of microbial eukaryotes at the Blanes Bay (13-19% of the OTUs depending on the analyzed size fraction, and ca. 40% of the sequences). Besides the distinct nature of prokaryotes and eukaryotes, this disparity could be explained by the differences in the data analysis, since Giner et al. (2019) used 99% clustering OTUs instead of ASVs and quantified recurrence using a metric developed of their own. Nevertheless, the number of seasonal ASVs we observed in bacteria triplicates the results found by Lambert et al. (2018) (89 ASVs), and the total relative abundance of seasonal organisms was also higher in our study compared to that observed at the SOLA station (47% vs 31.3%). Since we followed identical statistical methodologies and there is relatively high similarity between the environmental parameters and the number of ASVs, the observed differences were somehow surprising. A possible explanation could be related to the amplicon resolution, since for bacteria, Lambert et al. (2018) reported sequencing problems for the reverse complement pairs of Illumina sequencing (R2), analyzing thus 300 nucleotides instead of 490 as in here. Yet, this could explain a coarser taxonomy but not the changes in the total relative abundances of seasonal ASVs. The length of the time series was similar (7 years vs 11 years) and the sampling scheme, with biweekly sampling, could result to a certain degree in the disparities observed. Another explanation could derive from the presence of more irregular river discharges in the Banyuls basin, affecting the recurrence of the community through more variable salinity levels (Guizien et al., 2007). In any case, further studies would be needed to find a possible explanation for this discrepancy.

The seasonal patterns observed in our time series varied between different taxonomic groups (Supplementary Figure 5). Pelagibacter_A (SAR11 clade II) did not present seasonal ASVs. This result contrasts with what was observed in the Bermuda Atlantic Time series (BATS), in which this group is present mostly during spring (Giovannoni, 2017). On the other hand, AG-337-I02 (order HIMB59) peaked during winter, coinciding with what was observed at BATS (using SAR11 clade V as the group nomenclature). Nevertheless, the biogeochemical setting, physical forcing and other environmental factors that could control the temporal dynamics at BATS (Steinberg et al., 2001) are quite different from those of the coastal NW Mediterranean. Besides, HIMB114 (SAR11 clade III) in our study presented peak abundances during summer, a result also observed in Banyuls-sur-Mer (Salter et al., 2015). Overall, the observed differences in seasonal patterns among different sites point to the need of a deeper exploration of the niche of these groups, to investigate whether these differences have an ecological meaning or are due to methodological aspects.

### Niche similarity decreases with genetic distance

A clear trend between niche similarity and nucleotide divergence was detected for *Pelagibacter*, Pelagibacter_A and SAR86A. All these groups (i.e. SAR11 clade I and II) are known to contain many species with streamlined genomes and oligotrophic lifestyles (Dupont et al., 2012; Giovannoni, 2017). The pattern observed within these groups is consistent with habitat filtering (selection), in which similar niches are occupied by the same or genetically similar taxa. This pattern has already been observed in other environments (Horner-Devine & Bohannan, 2006; Tromas et al., 2018) and, interestingly, in our study we only observed it for groups with small genomes, which could be more affected by niche specialization. It is in fact unclear if closely related taxa compete. The evolution and diversification of traits between closely related taxa would allow their coexistence maintaining simultaneously the same realized niche (Martiny et al. 2015) as it was observed in our study for certain taxa. Trait diversification could arise from horizontal gene transfer events creating a larger pangenome for the different ecotypes. This in fact has been shown for *Pelagibacter*, from which the distinct genomes conforming its pangenome present differences in accessory genes between ecotypes with a 99.4% 16S rRNA gene identity (which corresponds to ~3 nucleotide divergences in our study; Delmont et al. 2019). Actually, we only detected a niche similarity pattern for *Pelagibacter*, Pelagibacter_A and SAR86A. Yet, we could have missed it for other genera due to lack of statistical power associated with sequencing depth. Thus, in order to determine if the niche similarity pattern is a common trait for all genera, we aggregated the non-significant values of all other genera detected in our study (details not shown) resulting in a non-significant pattern, and therefore, based on our data, we cannot conclude that this is a common pattern for marine bacteria. Nevertheless, we were able to describe how niche preference changes in relation to phylogenetic distance for three relevant marine groups. To dig further into the patterns of other groups, deeper sequencing or the sequencing of a larger 16SrRNA gene fragment is needed in order to improve the resolution and the number of variants obtained (Callahan et al., 2019, 2020).

When we checked how the individual ASVs responded to the measured environmental variables, we found two types of responses at the genus level: groups where all the ASVs displayed a similar response, such as *Pelagibacter*, AG-337-I02 (AEGEAN-169), D2472 (SAR86) and *Luminiphilus*, and groups with ASVs presenting niche differentiation, such as *Synechococcus* and SAR86A. The groups presenting the same patterns varied in the response; in the case of *Pelagibacter*, there was a clear distinction between the seasonal ecotypes and the ones appearing all year round (e.g. in Figure 4, the *Pelagibacter* dendrogram presents two clusters). The genera with distinct responses showed ecotype differentiation through niche partitioning processes. As an example, *Synechococcus* included ASVs with a positive response to temperature and other parameters, and ASVs with the opposite trend. Between the *Synechococcus,* ASV1 and ASV5, there were only 3-nucleotide divergence (99.26% identity), but the niche was clearly partitioned (Supplementary Figure 11). Since *Synechococcus* is one of the best known picoplankton groups, we checked the taxonomy at a finer resolution using a picocyanobacterial-specific database (Garczarek et al., 2020). In particular, ASV5 presented a 100% identity match with strain PROS-9-1, belonging to clade Ib, found in cold or temperate waters (Farrant et al., 2016). ASV1, on the other hand, resulted in a 100% match with members from multiple clades (clade I, II and III). This multiple match is an example of the problems with the limited power of the 16S rRNA gene V3-V4 region to resolve species (Johnson et al., 2019), with multiple clades possibly conforming the same ASVs, which could be an explanation to the dominant whole-year abundance of this variant. In our long-term dataset, we found that the peaks of ASV5 correspond to the recurrent yet temporall restricted *Synechococcus* bloom observed during spring with flow cytometry (Supplementary Figure 11). Summing up, these results illustrate the diversity of ecological trends within each genus. *Pelagibacter* ASVs presented similar ecological patterns, while other groups such as SAR86, HIMB59 and Synechococcus_C presented a clear ecotype differentiation. These within-genera differences would be hidden using clustering thresholds or working directly with the aggregation at the OTU 99% or genus level. Instead, our threshold-free analyses allowed to differentiate the responses at the ASV level, showing how there are taxa within the same genus presenting differentiated seasonality patterns even among closely related ASVs.

### Lack of seasonality at the class level

It has been hypothesized that bacteria from the same genus, family, order or even class could share ecological traits and respond similarly to environmental changes (Martiny et al., 2015; Philippot et al., 2010). In fact, it is unclear whether phylogenetic ranks are ecologically cohesive, and if true, to what rank this cohesiveness is maintained (Philippot et al., 2010). These ecological traits could be clearly determined by phylogenetic history, as is the case of particle versus free living lifestyle observed in deep ocean waters (Salazar et al., 2015). In the case of surface coastal waters, the periodic changes in environmental conditions should promote recurrent niches. We checked how seasonality was taxonomically clustered through testing the peak normalized power (PN) and its significance at various phylogenetic levels. By randomly aggregating the ASVs at different ranks, broad patterns of abundance could emerge coming from cohesive seasonal responses. When we tested whether this was true, we observed: a) groups that were always non-seasonal, b) groups with mixed responses with both seasonal and non-seasonal members, and c) groups that were always seasonal. The non-seasonal groups arise either from lack of seasonality signal or from multiple unsynchronized seasonal signals that generated a random and weak global signal. This was the case of the analyzed class rank levels, with all the results being non-seasonal (Figure 5). This seasonality and recurrence was opposite to that observed in the English Channel, with the Alphaproteobacteria and Gammaproteobacteria classes presenting a high autocorrelation and, therefore, a strong seasonal pattern (Faust et al., 2015; Gilbert et al., 2012). A possible explanation to these differences is that the English Channel presents much higher annual variability and a higher temperature range than Blanes Bay, therefore likely producing stronger habitat filtering. Bimodal distributions (seasonal and non-seasonal results) originate in groups containing ASVs that have strong seasonal trends and other non-seasonal ASVs, as is the case for Rhodobacterales and Pseudomonadales, copiotrophic groups occupying many different ecologic niches. *Rhodobacteraceae*, for example, includes ASVs with seasonality peaks in every season (Supplementary Figure 5). Finally, the seasonal groups were composed mostly by seasonal ASVs with most or all of them sharing the same time of the peak. The groups with all ASVs being seasonal could present more constrained optimal conditions of growth than the groups that appear randomly or all year-round. Examples of this behavior are the Puniceispirillales (SAR116 clade), a group harboring proteorhodopsin (Lee et al., 2019) and with most of the ASVs being seasonal and peaking during summer (Lee et al., 2019). Metagenomic and genome-centric approaches as well as physiological experimentation with available isolates would help shedding some light on the traits that determine the niche for these cohesive groups and the differences with other more diverse groups.

## Conclusions

The use of long-term time series and fine resolution of biological units allowed to compare within-genus ecological distributions. Specifically, we could prove that for certain genera niche similarity decreased with nucleotide divergence, indicating that multiple variants coexist due to habitat filtering processes. Additionally, through modeling of the differential abundance with a variety of environmental parameters, we unveiled some cases of niche partitioning resulting in different ecotypes producing blooms at different seasons. Finally, the analysis of different seasonality distributions for each phylogenetic rank (class, order, family, genus) indicated that the class rank was always non-seasonal for the groups analyzed, and thus ecologically non-coherent. This study sheds light into the niche specialization of various of the predominant genera in marine coastal microbial communities.

## Material and methods

### Location and sample collection

Samples were collected from the Blanes Bay Microbial Observatory, a station located in the NW Mediterranean sea about 1 km offshore over a water column of 20 m depth (41°40’N, 2°48’E; Gasol et al. 2016). Sampling was conducted monthly over 11 years, from January 2003 to December 2013. Water temperature and salinity were measured *in situ* with a conductivity, temperature and depth probe, and light penetration was estimated using a Secchi disk. Surface seawater was pre-filtered through a 200 μm nylon mesh, transported to the laboratory under dim light in 20 L plastic carboys, and processed within 2 h. Chlorophyll *a* concentration was measured on GF/F filters extracted with acetone and processed by fluorometry (Yentsch & Menzel, 1963). The concentrations of inorganic nutrients (NO_3_^−^, NO_2_^−^, NH_4_^+^, PO_4_^3-^, SiO_2_) were determined spectrophotometrically using an Alliance Evolution II autoanalyzer (Grasshoff et al., 1983). The abundances of picocyanobacteria, heterotrophic bacteria and photosynthetic pico- and nanoeukaryotes were determined by flow cytometry as described elsewhere (Gasol & Morán, 2016). Additionally, the abundance of photosynthetic and heterotrophic flagellates of different size ranges were measured by epifluorescence microscopy of filtrates on 0.6 μm polycarbonate filters stained with 4ʹ,6-diamidino-2-phenylindole. Microbial biomass was collected by filtering about 4 L of seawater using a peristaltic pump sequentially through a 20 μm nylon mesh (to remove large eukaryotes), a 3 μm pore-size 47 mm polycarbonate filter and a 0.2 μm pore-size Sterivex unit (Millipore).

### DNA extraction, PCR amplification and sequencing

DNA was extracted from the Sterivex unit with lysozyme, proteinase K and sodium dodecyl sulfate, and a standard phenol-chloroform-isoamyl alcohol protocol as described in Massana et al. (1997). The DNA analyzed here corresponds to the 0.2 to 3 μm fraction of bacterioplankton. Extracted DNA was purified and concentrated in an Amicon 100 (Millipore) and quantified in a NanoDrop-1000 spectrophotometer (Thermo Scientific). DNA was stored at −80⍛C and an aliquot from each sample was sent for sequencing to the Research and Testing Laboratory (Lubbock, TX, USA; http://rtlgenomics.com/). Primers 341F (5’-CCTACGGGNGGCWGCAG-3’, Herlemann et al. 2011) and 806RB (5’-GGACTACNVGGGTWTCTAAT-3’, Apprill et al. 2015) were used to amplify the V3-V4 regions of the 16S rRNA gene. A total of 131 samples were successfully sequenced and used in subsequent analyses.

### Sequence processing

*DADA2* v1.12 was used to differentiate the partial 16S rRNA gene amplicon sequence variants (ASVs) and to remove chimeras (parameters: maxN = 0, maxEE = 2,4, trunclen = 230,225; Callahan et al., 2016). Previously, spurious sequences and primers were trimmed using *cutadapt* v.1.16 (default values; M. Martin 2011). Taxonomic assignment of the ASVs was performed with IDTAXA from *DECIPHER* v2.14 package (40 confidence, Wright 2016) against the Genome Taxonomy Database (GTDB) r89 (Parks et al., 2018). IDTAXA reduces over classification, since most contemporary taxonomical databases are far from comprehensive and often lead to the misclassification of new groups. The GTDB has the advantage that it incorporates new data from metagenomic assembled genomes (MAGs) and generates phylogenies based on 120 single copy genes, resulting in a more robust phylogenetic tree than that created using only a single marker gene. Additionally, SILVA r138 taxonomy was used for nomenclature correspondence (Quast et al. 2013; see the correspondence between databases in Supplementary Table 2). The use of GTDB allowed an increase of assignation at the genus level (14.6% more sequences reaching the genus rank assignation) and the differentiation of new groups (e.g. D2472 genus within SAR86). Furthermore, the ASVs assigned to *Synechococcus* were checked against the Cyanorak database v2.1 (Garczarek et al., 2020) through 100% BLAST matches. ASVs classified as Mitochondria or Chloroplast were removed. The ASV sequences were also clustered into OTUs (Operational Taxonomic Units) at 97 and 99% identity in order to compare seasonal patterns at different similarity levels. Clustering was performed aligning all sequences, calculating a nucleotide distance matrix and identifying the clusters through the complete linkage method –maximum nucleotide distance between pairs of ASVs– using the *DECIPHER* package (Wright, 2016). This nucleotide distance matrix was also used to calculate the nucleotide divergence between ASVs.

### Community data analyses

We performed all analyses with the R v3.5 language (R Core Team, 2014). To process the data, we used the *phyloseq* v1.26 and *tidyverse* v1.3 packages (McMurdie & Holmes, 2013; Wickham et al., 2019) and *ggplot2* v3.2 for all visualizations (Wickham, 2016).

We defined abundant taxa as those above a 1% relative abundance in at least one sample as in Campbell et al. (2011). On the contrary, an ASV always below that cutoff was considered permanently rare. From both abundance groups, we defined three categories of ASVs based on their occurrence: broad (>75% occurrence), intermediate (>10% and <75% samples) and narrow (<10% samples) distributions, as termed by Chafee et al. (2018). The abundant ASVs were further tested as Conditionally Rare Taxa (CRT) –taxa typically in low abundance that occasionally become prevalent (bimodality =0.9, relative abundance threshold ≥0.5%)– following the description of Shade et al., (2014). The protocols test if each ASV follows a bimodal abundance distribution and if the values are above a minimum abundance threshold.

To estimate alpha diversity and beta diversity we used the *breakaway* v4.6 and *divnet* v0.34 packages respectively (A. Willis et al., 2017; A. D. Willis & Martin, 2020). These approaches avoid common pitfalls from applying classical ecology indexes (i.e. Chao1, Shannon, etc.) to microbiome data, which do not consider characteristics such as the influence of library size and compositionality.

### Seasonality data analysis

For seasonal analyses, the data was considered both at the month and season level, using for the latter the astronomical season definition as a delineation. To test whether each of the ASVs displayed seasonality –that is, recurrent changes over time– we used the lomb scargle periodogram (LSP) as implemented in the *lomb* package v1.2 (Ruf, 1999). This specific method accounts for unevenly sampled signals, a typical problem with long-term analyses. The method has already been used for testing the seasonality of marine microbial communities (see Lambert et al., 2018). Briefly, the LSP determines the spectrum of frequencies (the different sine waves with periods, for example half a year or one year) composing the dataset. Afterwards, through data randomizations, it tests whether the observed periods could occur by chance through a random distribution (*q* ≤ 0.01, FDR correction). For each ASV, we obtained the density distribution for each of the periods (a periodogram) and the peak normalized power (PN). The distribution shows which is the most recurrent period and the PN value measures the strength of this period. We followed the same criteria than Lambert et al., (2018) considering the results as seasonal only if PN was above 10 and *q* ≤ 0.01 (Lambert et al., 2018). We only examined ASVs present in at least 5% of the dataset (i.e. in at least 7 samples), resulting in 873 ASVs (corresponding to 94% of the total read relative abundance). In addition to the ASV level, we evaluated the seasonality at the class, order, family and genus taxonomic ranks. For a specific rank level (e.g. class Alphaproteobacteria), 80% of the ASVs conforming the group were chosen randomly, aggregated, and the LSP calculated. This process was repeated 300 times to obtain a distribution and observe how it compared to the LSP value without excluding any ASV. Out of the 29 classes present in the dataset, only the Alphaproteobacteria, Gammaproteobacteria and Bacteroidia could be evaluated since these are the classes that presented more than one order, family and genus ranks with at least 10 ASVs.

Further, we tested how the ASVs clustered based on the seasonal abundance patterns. We checked the number of possible clusters through the gap statistic from the *cluster* v2.1 package, since the expected number of clusters is unknown beforehand (Tibshirani et al., 2001). This approach tries to find the optimal *k* number of clusters by evaluating the drop of change between the normalized intra-clusters sum of squares distances (a measure of the compactness of the cluster, see Chapter 5 in Holmes and Huber, 2019). Once determined, we clustered the data through hierarchical clustering.

To visually compare the trend of the various seasonal ASVs, each one was fitted through a generalized additive model (GAM, *mgcv* v1.8 package, Hastie and Tibshirani 1986). A GAM is a generalized linear model in which the response variable depends linearly on various unknown smooth functions of some predictor variables. This method can fit polynomic responses without losing statistical relevance. The centered logarithm ratio values (pseudocount of 1) were fitted along the variable ‘day of the year’, allowing a smoothing parameter with 12 knots (the maximum number of curves to fit, being 12 for the number of months per year, Pedersen et al. 2019). Given the nature of the data (January evolves towards December and then the year starts again), a cyclic cubic spline condition was used to merge the start and end of the monthly distribution.

### Analyses of niche preference and environmental drivers

To examine how taxa within genus covary and, therefore, share a realized ecological niche, we used the *propr* v4.2 package (Quinn et al., 2017). This package was created to avoid the common pitfalls of compositional data analyzing correlation-like measurements. This particularity of our data creates many spurious correlations between the different taxa in which we cannot predict the true direction of change (i.e. in a community containing taxa A, B, C, is taxa A increasing or are taxa B and C decreasing? With relative abundances there is no distinction; see Gloor et al., 2017). A solution to this problem is to work with ratios instead of relative abundance. These ratios are usually obtained between the abundance of the taxon of interest and the geometric mean of all taxa for a specific sample (centered logarithm ratio, CLR). Then for all the ratios of taxa A and taxa B we measure the proportionality of change (Rho), which indicates how similar the abundance changes across many samples are. Two vectors (*x* and *y*) completely proportional (Rho=1) would present a variance of 0 for the ratio. The measure therefore presents similar properties to the correlation measurement (see Lovell et al. 2015 for a detailed explanation). The Rho statistic results were filtered with a final estimate of 5% of false discovery rate (FDR). Within each genus, we compared the Rho value between pairs of ASVs –acting as a proxy of niche similarity– against the nucleotide divergence among ASVs to see if there were trends in niche relatedness. A linear model was used to test which genera presented significant relationships (*p* < 0.05) between nucleotide divergence and Rho. We analyzed the genera with at least 10 closely related ASVs (at a maximum of 5 nucleotide divergence) which resulted in a total of 8 genera (out of 93). For most of these groups, using the V3 and V4 hypervariable regions of the 16S rRNA, 5 nucleotide divergence equals to a median sequence identity of 98.8% between two pairs. This nucleotide distance is the threshold that we use for considering two ASVs as closely related.

Finally, we tested which measured environmental parameters drive the patterns among closely related taxa. From the suite of measured variables, we selected temperature, total chlorophyll *a* concentration, inorganic nutrient concentrations, and the abundance of photosynthetic nanoflagellates (PNF) and heterotrophic nanoflagellates (HNF). Parameter selection was performed based on the expected relevance in modulating the ASV response (bottom up and top down processes) and also considering the number of missing values in the dataset. Multicollinearity between the parameters was tested using the *HH* v3.1 package (Heiberger, 2020). The variables presented a mean variance inflation factor (VIF) of 2. Only values of VIF exceeding 5 are considered as evidence of collinearity. To model the association we used the *corncob* v0.1 package (B. D. Martin et al., 2020), modeling each ASV across the different parameters and considering the values with an FDR ≤ 5% as significant. Afterwards, a display of the results was created with the GAM approach. The GAMs were applied to the data previously normalized through the centered logarithm ratio, using the geometric mean of the sample as denominator in the ratio (after adding a pseudocount of 1). Phosphate and nitrate concentrations, and the abundance of photosynthetic nanoflagellates displayed outliers in their distributions. The models were run with and without these values, generating similar results, and therefore we kept the outliers (details not shown).

### Reproducibility

The code for sequence data preprocessing, statistical analyses and visualization is available in the following repository: https://github.com/adriaaulaICM/bbmo_niche_sea. Sequence data have been deposited in the European Nucleotide Archive under project number PRJEB38773.

## Acknowledgments

We thank all the people involved in operating the BBMO, especially Clara Cardelús and Anselm for facilitating sampling, Vanessa Balagué for laboratory procedures and Ramon Massana for sharing the HNF data. We also thank the MarBits Bioinformatics platform of the Institut de Ciències del Mar, in particular Pablo Sánchez for computing support. We additionally thank Maria Touceda-Suárez, Gabriele Schiro and Yongjian Chen from the Barberán Lab for insightful discussions. This research was funded by grants REMEI (CTM2015-70340-R), MIAU (RTI2018-101025-B-I00) and ECLIPSE (PID2019-110128RB-I00) from the Spanish Ministry of Science and Innovation and Grup de Recerca de la Generalitat de Catalunya 2017SGR/1568. Adrià Auladell was supported by a Spanish FPI grant.

## Supplementary Table legends

**Supplementary Table 1:** Taxonomy and occurrence distribution of each individual ASV. ASV name, taxonomy (from domain to genus), presence (abundant or rare), distribution (broad, intermediate or narrow), Conditionally Rare taxa (CRT) and ASV seasonality.

**Supplementary Table 2:** Correspondence between the GTDB and SILVA genus nomenclature. The first two columns correspond to the genus, family and order from the GTDB r89, and the next two provide the same information in SILVA DB r138. N. seasonal indicates the number of seasonal ASVs from the total of ASVs tested. Finally, the column “General Information Genus” provides useful information behind some of the changes in the nomenclature.

**Supplementary Table 3:** Linear regression coefficients for each genus between Rho proportionality values and nucleotide divergence. Df, degrees of freedom; logLik, log likelihood; AIC, Akaike Information Criterion; BIC Bayesian Information Criterion; deviance; df.residual, residual degrees of freedom; pval.term, *p* values of the coefficient; R.square.

## References

Alonso-Sáez, L., Balagué, V., Sà, E. L., Sánchez, O., González, J. M., Pinhassi, J., Massana, R., Pernthaler, J., Pedrós-Alió, C., & Gasol, J. M. (2007). Seasonality in bacterial diversity in north-west Mediterranean coastal waters: Assessment through clone libraries, fingerprinting and FISH. FEMS Microbiology Ecology, 60(1), 98–112. https://doi.org/10.1111/j.1574-6941.2006.00276.x

Apprill, A., McNally, S., Parsons, R., & Weber, L. (2015). Minor revision to V4 region SSU rRNA 806R gene primer greatly increases detection of SAR11 bacterioplankton. Aquatic Microbial Ecology, 75(2), 129–137. https://doi.org/10.3354/ame01753

Buttigieg, P. L., Fadeev, E., Bienhold, C., Hehemann, L., Offre, P., & Boetius, A. (2018). Marine microbes in 4D — using time series observation to assess the dynamics of the ocean microbiome and its links to ocean health. Current Opinion in Microbiology, 43, 169–185. https://doi.org/10.1016/j.mib.2018.01.015

Callahan, B. J., Grinevich, D., Thakur, S., Balamotis, M. A., & Yehezkel, T. B. (2020). Ultra-accurate Microbial Amplicon Sequencing Directly from Complex Samples with Synthetic Long Reads. BioRxiv, 2020.07.07.192286. https://doi.org/10.1101/2020.07.07.192286

Callahan, B. J., McMurdie, P. J., & Holmes, S. P. (2017). Exact sequence variants should replace operational taxonomic units in marker-gene data analysis. The ISME Journal, 11(12), 2639–2643. https://doi.org/10.1038/ismej.2017.119

Callahan, B. J., McMurdie, P. J., Rosen, M. J., Han, A. W., Johnson, A. J. A., & Holmes, S. P. (2016). DADA2: High-resolution sample inference from Illumina amplicon data. Nature Methods, 13, 581.

Callahan, B. J., Wong, J., Heiner, C., Oh, S., Theriot, C. M., Gulati, A. S., McGill, S. K., & Dougherty, M. K. (2019). High-throughput amplicon sequencing of the full-length 16S rRNA gene with single-nucleotide resolution. Nucleic Acids Research, 47(18), e103–e103. https://doi.org/10.1093/nar/gkz569

Campbell, B. J., Yu, L., Heidelberg, J. F., & Kirchman, D. L. (2011). Activity of abundant and rare bacteria in a coastal ocean. Proceedings of the National Academy of Sciences, 108(31), 12776–12781. https://doi.org/10.1073/pnas.1101405108

Chafee, M., Fernàndez-Guerra, A., Buttigieg, P. L., Gerdts, G., Eren, A. M., Teeling, H., & Amann, R. I. (2018). Recurrent patterns of microdiversity in a temperate coastal marine environment. ISME Journal, 12(1), 237–252. https://doi.org/10.1038/ismej.2017.165

Chow, C.-E. T., Sachdeva, R., Cram, J. A., Steele, J. A., Needham, D. M., Patel, A., Parada, A. E., & Fuhrman, J. A. (2013). Temporal variability and coherence of euphotic zone bacterial communities over a decade in the Southern California Bight. The ISME Journal, 7(12), 2259–2273. https://doi.org/10.1038/ismej.2013.122

Cohan, F. M. (2017). Transmission in the Origins of Bacterial Diversity, From Ecotypes to Phyla. Microbiology Spectrum, 5(5). https://doi.org/10.1128/microbiolspec.MTBP-0014-2016

Cram, J. A., Chow, C.-E. T., Sachdeva, R., Needham, D. M., Parada, A. E., Steele, J. A., & Fuhrman, J. A. (2015). Seasonal and interannual variability of the marine bacterioplankton community throughout the water column over ten years. The ISME Journal, 9(3), 563–580. https://doi.org/10.1038/ismej.2014.153

Delmont, T. O., Kiefl, E., Kilinc, O., Esen, O. C., Uysal, I., Rappé, M. S., Giovannoni, S., & Eren, A. M. (2019). Single-amino acid variants reveal evolutionary processes that shape the biogeography of a global SAR11 subclade. ELife, 8, e46497. https://doi.org/10.7554/eLife.46497

Dupont, C. L., Rusch, D. B., Yooseph, S., Lombardo, M.-J., Alexander Richter, R., Valas, R., Novotny, M., Yee-Greenbaum, J., Selengut, J. D., Haft, D. H., Halpern, A. L., Lasken, R. S., Nealson, K., Friedman, R., & Craig Venter, J. (2012). Genomic insights to SAR86, an abundant and uncultivated marine bacterial lineage. The ISME Journal, 6(6), 1186–1199. https://doi.org/10.1038/ismej.2011.189

Eren, A. M., Maignien, L., Sul, W. J., Murphy, L. G., Grim, S. L., Morrison, H. G., & Sogin, M. L. (2013). Oligotyping: Differentiating between closely related microbial taxa using 16S rRNA gene data. Methods in Ecology and Evolution, 4(12), 1111–1119. https://doi.org/10.1111/2041-210X.12114

Falkowski, P. (2012). Ocean Science: The power of plankton. Nature, 483(7387), S17–S20. https://doi.org/10.1038/483S17a

Farrant, G. K., Doré, H., Cornejo-Castillo, F. M., Partensky, F., Ratin, M., Ostrowski, M., Pitt, F. D., Wincker, P., Scanlan, D. J., Iudicone, D., Acinas, S. G., & Garczarek, L. (2016). Delineating ecologically significant taxonomic units from global patterns of marine picocyanobacteria. Proceedings of the National Academy of Sciences, 113(24), E3365–E3374. https://doi.org/10.1073/pnas.1524865113

Faust, K., Lahti, L., Gonze, D., de Vos, W. M., & Raes, J. (2015). Metagenomics meets time series analysis: Unraveling microbial community dynamics. Current Opinion in Microbiology, 25(May), 56–66. https://doi.org/10.1016/j.mib.2015.04.004

Friedman, J., & Alm, E. J. (2012). Inferring Correlation Networks from Genomic Survey Data. PLoS Computational Biology, 8(9), e1002687. https://doi.org/10.1371/journal.pcbi.1002687

Fuhrman, J. A., Cram, J. A., & Needham, D. M. (2015). Marine microbial community dynamics and their ecological interpretation. Nature Reviews: Microbiology, 13(3), 133–146. https://doi.org/10.1038/nrmicro3417

Garczarek, L., Guyet, U., Doré, H., Farrant, G. K., Hoebeke, M., Brillet-Guéguen, L., Bisch, A., Ferrieux, M., Siltanen, J., Corre, E., Le Corguillé, G., Ratin, M., Pitt, F. D., Ostrowski, M., Conan, M., Siegel, A., Labadie, K., Aury, J.-M., Wincker, P., … Partensky, F. (2020). Cyanorak v2.1: A scalable information system dedicated to the visualization and expert curation of marine and brackish picocyanobacteria genomes. Nucleic Acids Research, gkaa958. https://doi.org/10.1093/nar/gkaa958

Gasol, J. M., Cardelús, C., Morán, X. A. G., Balagué, V., Forn, I., Marrasé, C., Massana, R., Pedrós-Alió, C., Sala, M. M., Simó, R., Vaqué, D., & Estrada, M. (2016). Seasonal patterns in phytoplankton photosynthetic parameters and primary production at a coastal NW Mediterranean site. Scientia Marina, 80S1, 63–77. http://dx.doi.org/10.3989/scimar.04480.06E

Gasol, J. M., & Morán, X. A. G. (2016). Flow Cytometric Determination of Microbial Abundances and Its Use to Obtain Indices of Community Structure and Relative Activity. In T. J. McGenity, K. N. Timmis, & B. Nogales (Eds.), Hydrocarbon and Lipid Microbiology Protocols: Single-Cell and Single-Molecule Methods (pp. 159–187). Springer. https://doi.org/10.1007/8623_2015_139

Ghai, R., Mizuno, C. M., Picazo, A., Camacho, A., & Rodriguez-Valera, F. (2013). Metagenomics uncovers a new group of low GC and ultra-small marine Actinobacteria. Scientific Reports, 3(1), 1–8. https://doi.org/10.1038/srep02471

Gilbert, J. A., Steele, J. A., Caporaso, J. G., Steinbrück, L., Reeder, J., Temperton, B., Huse, S., McHardy, A. C., Knight, R., Joint, I., Somerfield, P., Fuhrman, J. A., & Field, D. (2012). Defining seasonal marine microbial community dynamics. The ISME Journal, 6(2), 298–308. https://doi.org/10.1038/ismej.2011.107

Giner, C. R., Balagué, V., Krabberød, A. K., Ferrera, I., Reñé, A., Garcés, E., Gasol, J. M., Logares, R., & Massana, R. (2019). Quantifying long-term recurrence in planktonic microbial eukaryotes. Molecular Ecology, 28(5), 923–935. https://doi.org/10.1111/mec.14929

Giovannoni, S. J. (2017). SAR11 Bacteria: The Most Abundant Plankton in the Oceans. Annual Review of Marine Science, 9(1), 231–255. https://doi.org/10.1146/annurev-marine-010814-015934

Gloor, G. B., Macklaim, J. M., Pawlowsky-Glahn, V., & Egozcue, J. J. (2017). Microbiome datasets are compositional: And this is not optional. Frontiers in Microbiology, 8(NOV), 1–6. https://doi.org/10.3389/fmicb.2017.02224

Grasshoff, K., Ehrhardt, M., & Kremling, K. (1983). Methods of seawater analysis (2nd ed.).

Guizien, K., Charles, F., Lantoine, F., & Naudin, J.-J. (2007). Nearshore dynamics of nutrients and chlorophyll during Mediterranean-type flash-floods. Aquatic Living Resources, 20(1), 3–14. https://doi.org/10.1051/alr:2007011

Hastie, T., & Tibshirani, R. (1986). Generalized Additive Models. Statistical Science, 1(3), 297–310. https://doi.org/10.1214/ss/1177013604

Heiberger, R. M. (2020). HH: Statistical Analysis and Data Display: Heiberger and Holland. (3.1–40) [Computer software]. https://CRAN.R-project.org/package=HH

Herlemann, D. P., Labrenz, M., Jürgens, K., Bertilsson, S., Waniek, J. J., & Andersson, A. F. (2011). Transitions in bacterial communities along the 2000 km salinity gradient of the Baltic Sea. The ISME Journal, 5(10), 1571–1579. https://doi.org/10.1038/ismej.2011.41

Holmes, S., & Huber, W. (2019). Modern Statistics for Modern Biology (1 edition). Cambridge University Press.

Horner-Devine, M. C., & Bohannan, B. J. M. (2006). PHYLOGENETIC CLUSTERING AND OVERDISPERSION IN BACTERIAL COMMUNITIES. Ecology, 87(sp7), S100–S108. https://doi.org/10.1890/0012-9658(2006)87[100:PCAOIB]2.0.CO;2

Hutchinson, G. E. (1957). Concluding Remarks. Cold Spring Harbor Symposia on Quantitative Biology, 22(0), 415–427. https://doi.org/10.1101/SQB.1957.022.01.039

Johnson, J. S., Spakowicz, D. J., Hong, B.-Y., Petersen, L. M., Demkowicz, P., Chen, L., Leopold, S. R., Hanson, B. M., Agresta, H. O., Gerstein, M., Sodergren, E., & Weinstock, G. M. (2019). Evaluation of 16S rRNA gene sequencing for species and strain-level microbiome analysis. Nature Communications, 10(1), 5029. https://doi.org/10.1038/s41467-019-13036-1

Ladau, J., & Eloe-Fadrosh, E. A. (2019). Spatial, Temporal, and Phylogenetic Scales of Microbial Ecology. Trends in Microbiology, 0(0). https://doi.org/10.1016/j.tim.2019.03.003

Lambert, S., Tragin, M., Lozano, J.-C., Ghiglione, J.-F., Vaulot, D., Bouget, F.-Y., & Galand, P. E. (2018). Rhythmicity of coastal marine picoeukaryotes, bacteria and archaea despite irregular environmental perturbations. The ISME Journal. https://doi.org/10.1038/s41396-018-0281-z

Lee, J., Kwon, K. K., Lim, S.-I., Song, J., Choi, A. R., Yang, S.-H., Jung, K.-H., Lee, J.-H., Kang, S. G., Oh, H.-M., & Cho, J.-C. (2019). Isolation, cultivation, and genome analysis of proteorhodopsin-containing SAR116-clade strain Candidatus Puniceispirillum marinum IMCC1322. Journal of Microbiology, 57(8), 676–687. https://doi.org/10.1007/s12275-019-9001-2

Logares, R., Deutschmann, I. M., Junger, P. C., Giner, C. R., Krabberød, A. K., Schmidt, T. S. B., Rubinat-Ripoll, L., Mestre, M., Salazar, G., Ruiz-González, C., Sebastián, M., de Vargas, C., Acinas, S. G., Duarte, C. M., Gasol, J. M., & Massana, R. (2020). Disentangling the mechanisms shaping the surface ocean microbiota. Microbiome, 8(1), 55. https://doi.org/10.1186/s40168-020-00827-8

Lovell, D., Pawlowsky-Glahn, V., Egozcue, J. J., Marguerat, S., & Bähler, J. (2015). Proportionality: A Valid Alternative to Correlation for Relative Data. PLoS Computational Biology, 11(3), e1004075. https://doi.org/10.1371/journal.pcbi.1004075

Martijn, J., Vosseberg, J., Guy, L., Offre, P., & Ettema, T. J. G. (2018). Deep mitochondrial origin outside the sampled alphaproteobacteria. Nature, 557(7703), 101–105. https://doi.org/10.1038/s41586-018-0059-5

Martin, B. D., Witten, D., & Willis, A. D. (2020). Modeling microbial abundances and dysbiosis with beta-binomial regression. Annals of Applied Statistics, 14(1), 94–115. https://doi.org/10.1214/19-AOAS1283

Martin, M. (2011). Cutadapt removes adapter sequences from high-throughput sequencing reads. EMBnet.Journal, 17(1), 10. https://doi.org/10.14806/ej.17.1.200

Martiny, J. B. H., Jones, S. E., Lennon, J. T., & Martiny, A. C. (2015). Microbiomes in light of traits: A phylogenetic perspective. Science, 350(6261). https://doi.org/10.1126/science.aac9323

Massana, R., Murray, A. E., Preston, C. M., & Delong, E. F. (1997). Vertical distribution and phylogenetic characterization of marine planktonic Archaea in the Santa Barbara Channel. Applied and Environmental Microbiology, 63(1), 50–56.

McMurdie, P. J., & Holmes, S. (2013). Phyloseq: An R package for reproducible interactive analysis and graphics of microbiome census data. PLoS ONE, 8(4). https://doi.org/10.1371/journal.pone.0061217

Mestre, M., Borrull, E., Sala, M. M., & Gasol, J. M. (2017). Patterns of bacterial diversity in the marine planktonic particulate matter continuum. The ISME Journal, 11(4), 999–1010. https://doi.org/10.1038/ismej.2016.166

Needham, D. M., Fichot, E. B., Wang, E., Berdjeb, L., Cram, J. A., Fichot, C. G., & Fuhrman, J. A. (2018). Dynamics and interactions of highly resolved marine plankton via automated high-frequency sampling. The ISME Journal, 12(10), 2417–2432. https://doi.org/10.1038/s41396-018-0169-y

Nunes, S., Latasa, M., Gasol, J. M., & Estrada, M. (2018). Seasonal and interannual variability of phytoplankton community structure in a Mediterranean coastal site. Marine Ecology Progress Series, 592, 57–75. https://doi.org/10.3354/meps12493

Parks, D. H., Chuvochina, M., Waite, D. W., Rinke, C., Skarshewski, A., Chaumeil, P.-A., & Hugenholtz, P. (2018). A standardized bacterial taxonomy based on genome phylogeny substantially revises the tree of life. Nature Biotechnology, 36(10), 996–1004. https://doi.org/10.1038/nbt.4229

Pedersen, E. J., Miller, D. L., Simpson, G. L., & Ross, N. (2019). Hierarchical generalized additive models in ecology: An introduction with mgcv. PeerJ, 7, e6876. https://doi.org/10.7717/peerj.6876

Philippot, L., Andersson, S. G. E., Battin, T. J., Prosser, J. I., Schimel, J. P., Whitman, W. B., & Hallin, S. (2010). The ecological coherence of high bacterial taxonomic ranks. Nature Reviews Microbiology, 8(7), 523–529. https://doi.org/10.1038/nrmicro2367

Quast, C., Pruesse, E., Yilmaz, P., Gerken, J., Schweer, T., Yarza, P., Peplies, J., & Glöckner, F. O. (2013). The SILVA ribosomal RNA gene database project: Improved data processing and web-based tools. Nucleic Acids Research, 41(Database issue), D590–D596. https://doi.org/10.1093/nar/gks1219

Quinn, T. P., Richardson, M. F., Lovell, D., & Crowley, T. M. (2017). propr: An R-package for Identifying Proportionally Abundant Features Using Compositional Data Analysis. Scientific Reports, 7(1), 1–9. https://doi.org/10.1038/s41598-017-16520-0

R Core Team. (2014). R: A language and environment for statistical computing. R Foundation for Statistical Computing, Vienna, Austria. https://www.r-project.org/

Ruf, T. (1999). The Lomb-Scargle Periodogram in Biological Rhythm Research: Analysis of Incomplete and Unequally Spaced Time-Series. Biological Rhythm Research, 30(2), 178–201. https://doi.org/10.1076/brhm.30.2.178.1422

Salazar, G., Cornejo-Castillo, F. M., Benítez-Barrios, V., Fraile-Nuez, E., Álvarez-Salgado, X. A., Duarte, C. M., Gasol, J. M., & Acinas, S. G. (2015). Global diversity and biogeography of deep-sea pelagic prokaryotes. The ISME Journal, 1–13. https://doi.org/10.1038/ismej.2015.137

Salter, I., Galand, P. E., Fagervold, S. K., Lebaron, P., Obernosterer, I., Oliver, M. J., Suzuki, M. T., & Tricoire, C. (2015). Seasonal dynamics of active SAR11 ecotypes in the oligotrophic Northwest Mediterranean Sea. The ISME Journal, 9(2), 347–360. https://doi.org/10.1038/ismej.2014.129

Shade, A., Jones, S. E., Caporaso, J. G., Handelsman, J., Knight, R., Fierer, N., & Gilbert, A. (2014). Conditionally rare taxa disproportionately contribute to temporal changes in microbial diversity. MBio, 5(4), 1–9. https://doi.org/10.1128/mBio.01371-14

Steinberg, D. K., Carlson, C. A., Bates, N. R., Johnson, R. J., Michaels, A. F., & Knap, A. H. (2001). Overview of the US JGOFS Bermuda Atlantic Time-series Study (BATS): A decade-scale look at ocean biology and biogeochemistry. Deep Sea Research Part II: Topical Studies in Oceanography, 48(8-9), 1405–1447. https://doi.org/10.1016/S0967-0645(00)00148-X

Sunagawa, S., Coelho, L. P., Chaffron, S., Kultima, J. R., Labadie, K., Salazar, G., Djahanschiri, B., Zeller, G., Mende, D. R., Alberti, A., Cornejo-Castillo, F. M., Costea, P. I., Cruaud, C., D’Ovidio, F., Engelen, S., Ferrera, I., Gasol, J. M., Guidi, L., Hildebrand, F., … Bork, P. (2015). Ocean plankton. Structure and function of the global ocean microbiome. Science, 348(6237), 1261359. https://doi.org/10.1126/science.1261359

Tibshirani, R., Walther, G., & Hastie, T. (2001). Estimating the number of clusters in a data set via the gap statistic. Journal of the Royal Statistical Society: Series C (Applied Statistics), 63(2), 411–423. https://doi.org/10.1111/1467-9868.00293

Tromas, N., Taranu, Z. E., Martin, B. D., Willis, A., Fortin, N., Greer, C. W., & Shapiro, B. J. (2018). Niche Separation Increases With Genetic Distance Among Bloom-Forming Cyanobacteria. Frontiers in Microbiology, 9, 438. https://doi.org/10.3389/fmicb.2018.00438

VanInsberghe, D., Arevalo, P., Chien, D., & Polz, M. F. (2020). How can microbial population genomics inform community ecology? Philosophical Transactions of the Royal Society B, 375(1798), 20190253. https://doi.org/10.1098/rstb.2019.0253

Viklund, J., Martijn, J., Ettema, T. J. G., & Andersson, S. G. E. (2013). Comparative and Phylogenomic Evidence That the Alphaproteobacterium HIMB59 Is Not a Member of the Oceanic SAR11 Clade. PLoS ONE, 8(11), e78858. https://doi.org/10.1371/journal.pone.0078858

Wickham, H. (2016). ggplot2: Elegant graphics for data analysis. Springer-Verlag New York. https://ggplot2.tidyverse.org

Wickham, H., Averick, M., Bryan, J., Chang, W., McGowan, L. D., François, R., Grolemund, G., Hayes, A., Henry, L., Hester, J., Kuhn, M., Pedersen, T. L., Miller, E., Bache, S. M., Müller, K., Ooms, J., Robinson, D., Seidel, D. P., Spinu, V., … Yutani, H. (2019). Welcome to the tidyverse. Journal of Open Source Software, 4(43), 1686. https://doi.org/10.21105/joss.01686

Willis, A., Bunge, J., & Whitman, T. (2017). Improved detection of changes in species richness in high diversity microbial communities. Journal of the Royal Statistical Society: Series C (Applied Statistics), 66(5), 963–977. https://doi.org/10.1111/rssc.12206

Willis, A. D., & Martin, B. D. (2020). Estimating diversity in networked ecological communities. Biostatistics, kxaa015. https://doi.org/10.1093/biostatistics/kxaa015

Wright, E., S. (2016). Using DECIPHER v2.0 to Analyze Big Biological Sequence Data in R. The R Journal, 8(1), 352. https://doi.org/10.32614/RJ-2016-025

Yentsch, C. S., & Menzel, D. W. (1963). A method for the determination of phytoplankton chlorophyll and phaeophytin by fluorescence. Deep Sea Research and Oceanographic Abstracts, 10(3), 221–231. https://doi.org/10.1016/0011-7471(63)90358-9

