## Supplementary figures (1-11) for "Seasonal niche differentiation between evolutionary closely related marine bacteria"

Adrià Auladell[1], Albert Barberán[2], Ramiro Logares[1], Esther Garcés[1], Josep M. Gasol[1,3], and Isabel Ferrera[1,4]

[1]Departament de Biologia Marina i Oceanografia, Institut de Ciències del Mar, ICM-CSIC, 08003 Barcelona, Catalunya, Spain
[2]Department of Environmental Science, University of Arizona, Tucson, 85721 AZ, USA
[3]Center for Marine Ecosystems Research, School of Science, Edith Cowan University, Joondalup, WA, Australia
[4]Centro Oceanográfico de Málaga, Instituto Español de Oceanografía, 29640 Fuengirola, Málaga, Spain

- - -

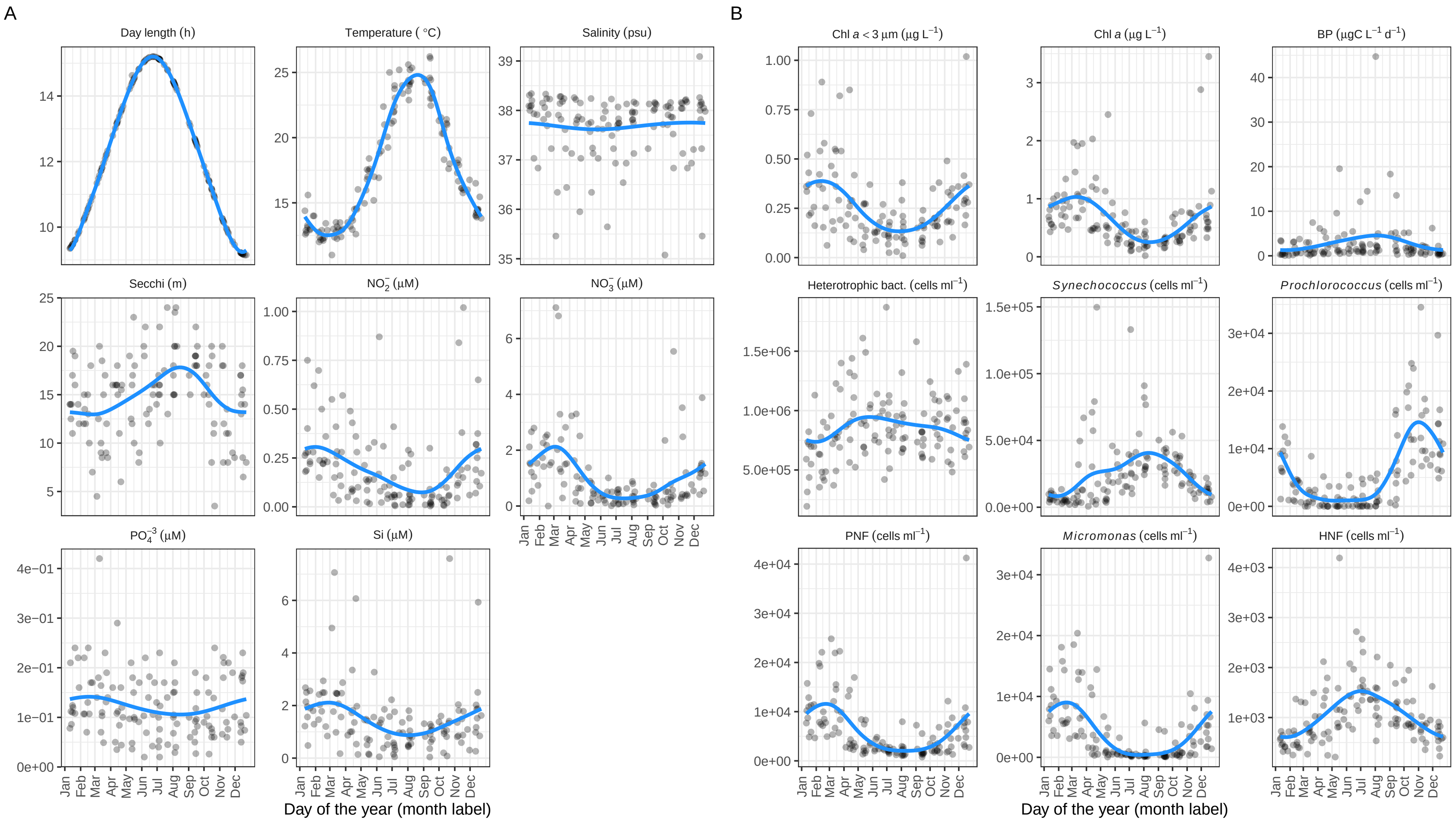

Figure S1: Distribution of the physicochemical (A) and biological (B) environmental variables measured in the Blanes Bay Microbial Observatory during the 11 years of this study. The Y axis corresponds to the parameter value (units indicated in the plot title) and the X axis corresponds to the day of the year (month is shown for orientation, with the line ticks for the first day). A generalized additive model was fitted to the data. BP: Bacterial production; PNF: Phototrophic nanoflagellates; Cha<3 $\mu$m: Chlorophyll *a* from the 3 $\mu$m fraction or smaller; HNF: Heterotrophic nanoflagellates.

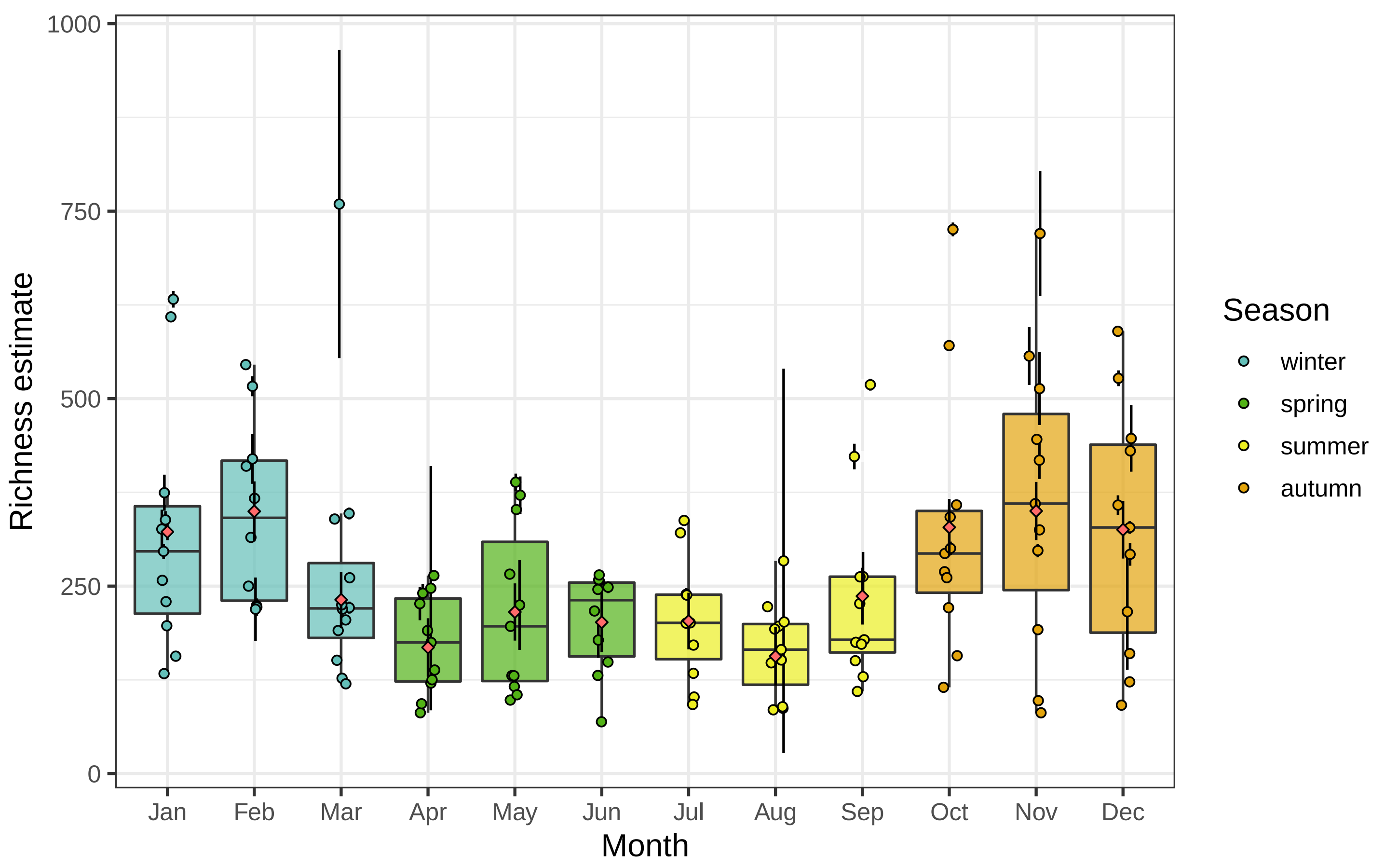

Figure S2: Alpha diversity (richness estimate) for the whole time series (11 years). Dots colored by season correspond to the sample estimates with the confidence interval at 95%; red rhomboids correspond to the mean month richness estimate by the *breakaway* package (with confidence interval 95%). Each boxplot presents the median and the 25% and 75% limits with the distribution of 11 points, and whiskers represent 1.5 times the interquartile range.

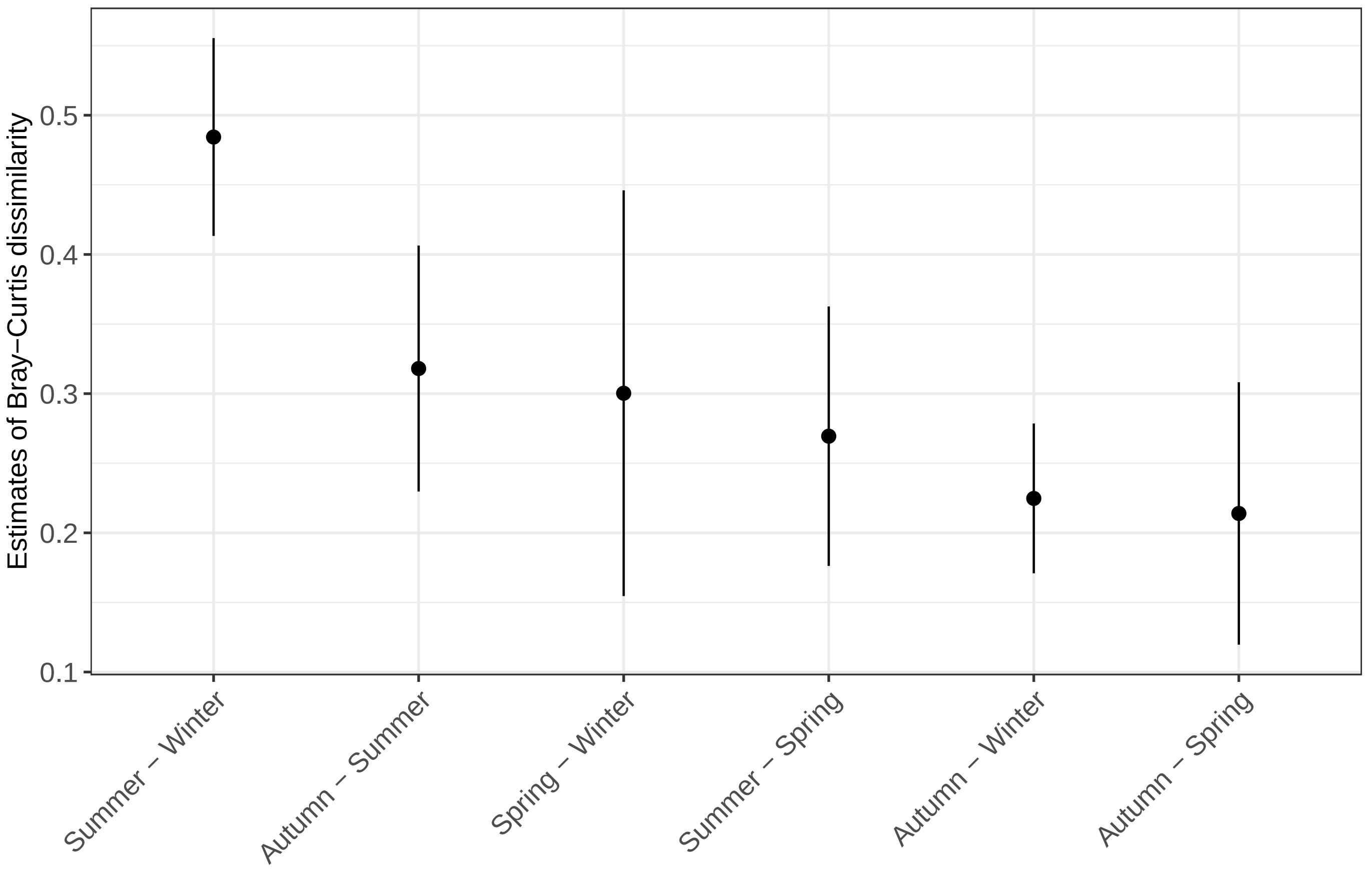

Figure S3: Comparison between the beta diversity estimates of Bray Curtis dissimilarity between the different seasons. The estimates and the 95% confidence intervals are displayed.

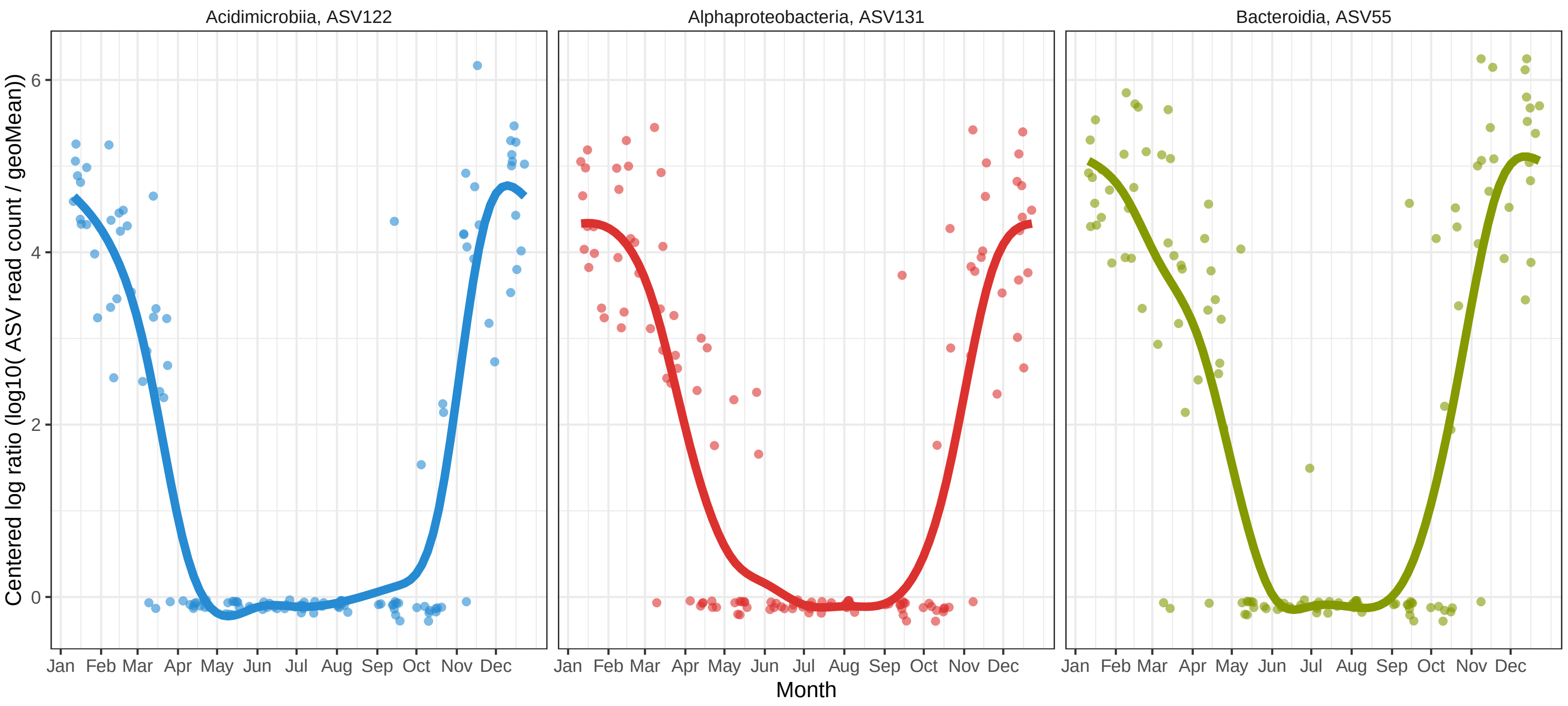

Figure S4: Distribution of some ASVs with autumn-winter seasonality. The X axis corresponds to the day of the year (month is shown for orientation, with the line ticks for the first day) and the Y axis presents the read count transformed through the centered logarithm ratio abundance. A generalized additive model smooth is adjusted to the data points. Taxonomic classification reached the class level only.

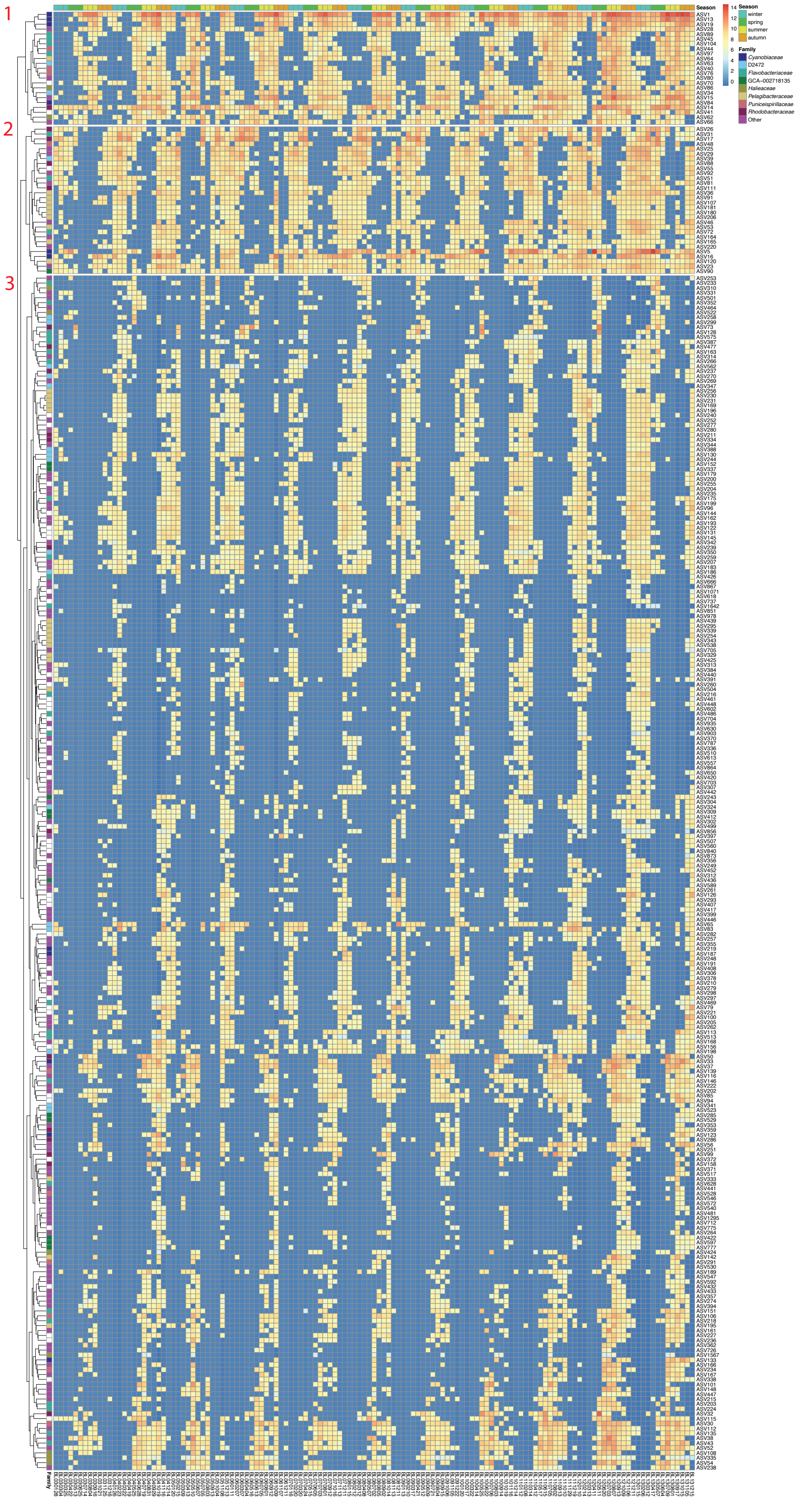

Figure S5: Heatmap displaying all the seasonal ASVs with a hierarchical clustering of the distributions. The 3 main clusters identified are presented and indicated with a red number. Each ASV is color coded by taxonomy and each sample with the season.

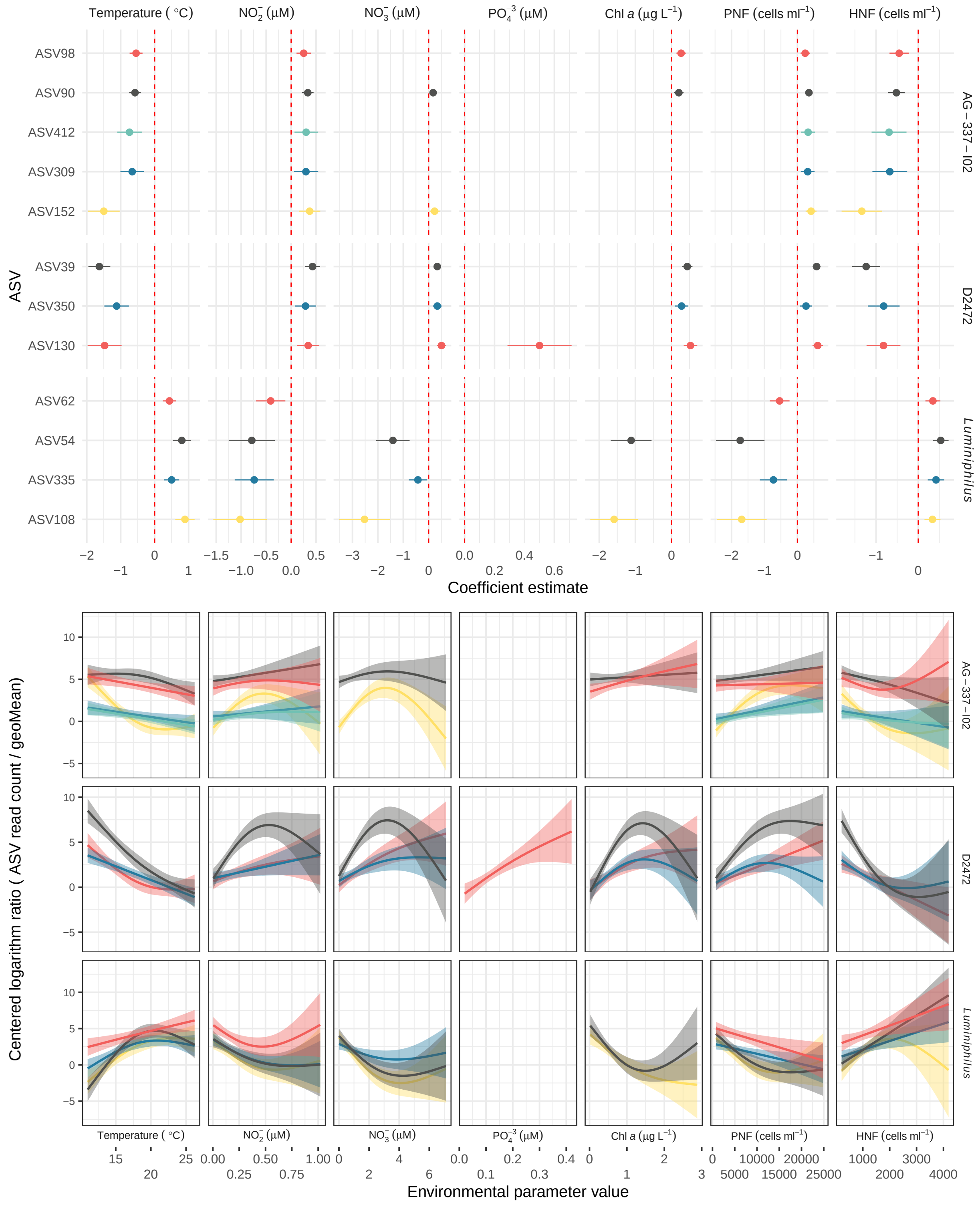

Figure S6: A) Significant models between ASVs from AG−337−I02 (Alphaproteobacteria), D2472 (Gammaproteobacteria) and *Luminiphilus* (Gammaproteobacteria) genera (rows) and environmental parameters (columns). The coefficient estimate indicates positive or negative responses to the parameter and is shown with a 95% confidence interval. The colors correspond to the different ASV within a genus (only top 8 abundant ASVs are colored, the other ASVs are shown in grey). ASVs are ordered through a hierarchical clustering based on nucleotide divergence. B) Generalized additive model fits between the ASV centered logarithm ratio abundances and the parameter value distribution for the significant ASVs indicated in the upper plot. Panels and ASV colors are distributed as in the upper panel. PNF: Phototrophic nanoflagelates; HNF: Heterotrophic nanoflagelates.

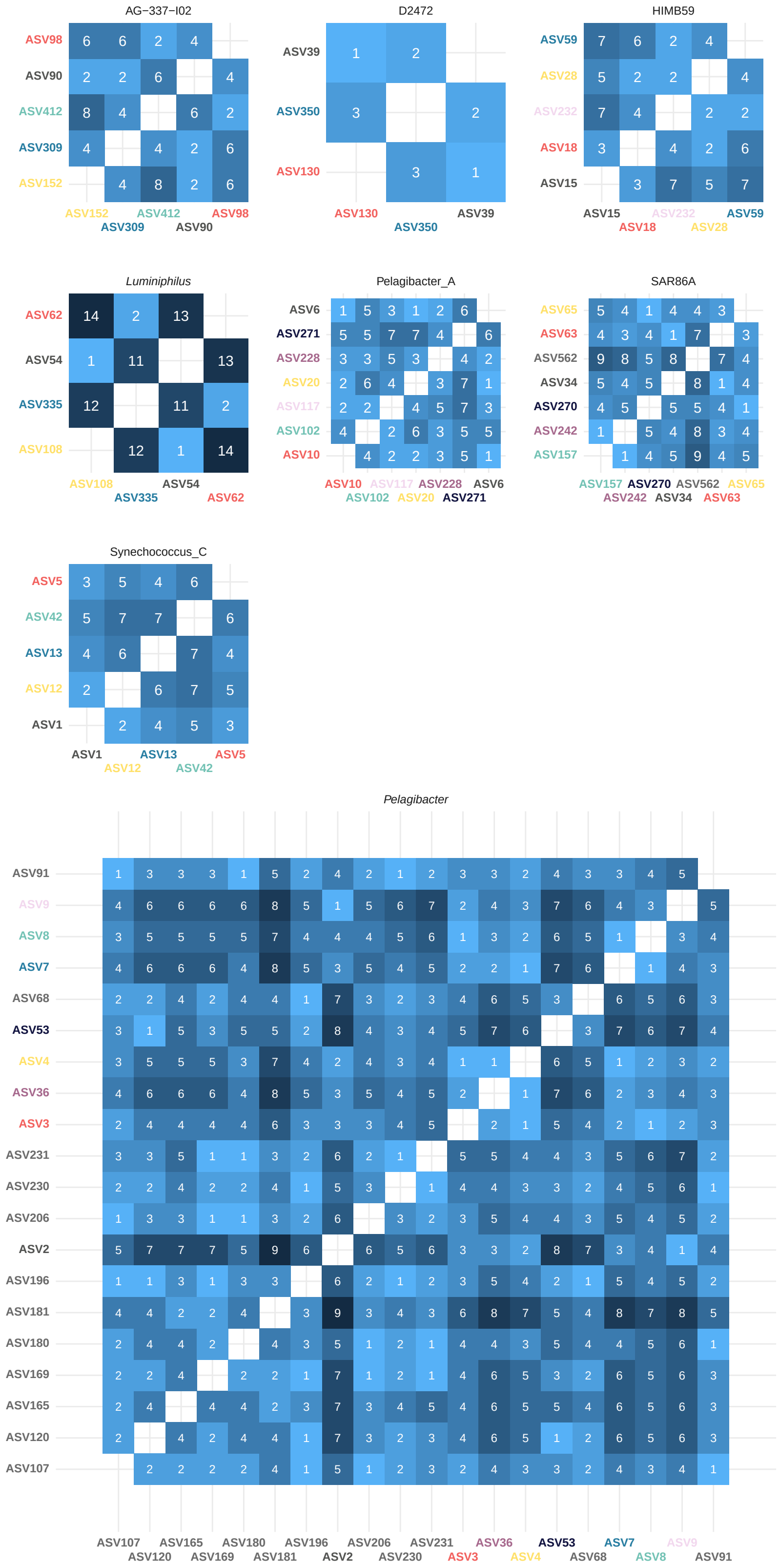

Figure S7: Nucleotide divergence heatmap between groups of ASVs presenting a significant response to environmental parameters. The genera included are AG−337−I02, HIMB59, Pelagibacter_A and *Pelagibacter* (Alphaproteobacteria); D2472, SAR86 and *Luminiphilus* (Gammaproteobacteria); and Synechococcus_C (Cyanobacteria). The color corresponds to the different ASVs within a genus (only the top 8 abundant ASVs are colored, the other ASVs are shown in grey). Five nucleotide divergence equals to a median sequence identity of 98.8%.

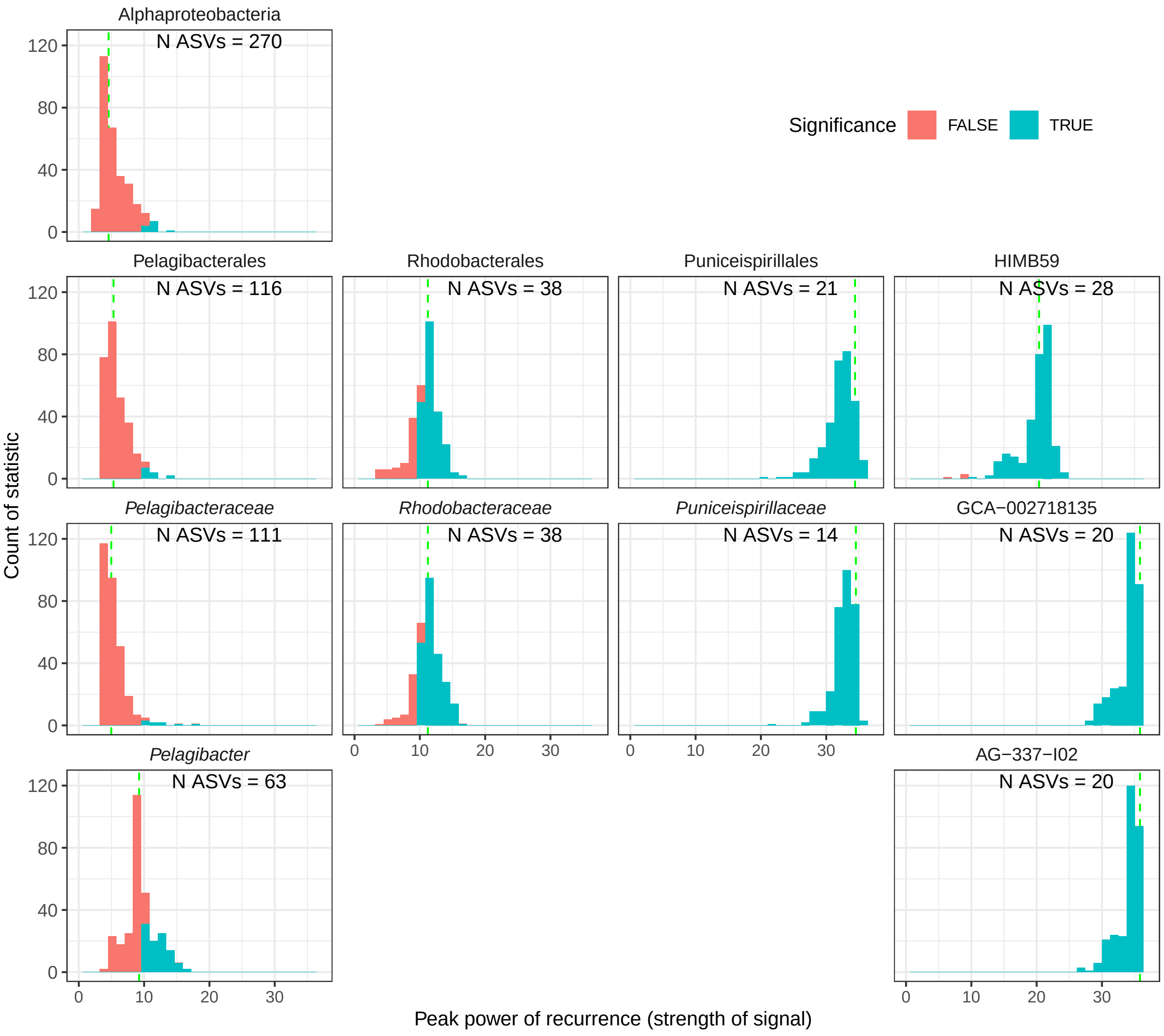

Figure S8: A) Histograms of the peak normalized power statistic at the class, order, family and genus level (from top to bottom, each line is a rank) of 80% of the ASVs conforming the rank for class Alphaproteobacteria. The red bins indicate the non-significant results (PN$\geq$10, $q\leq$0.01) and blue bins the significant ones. The dashed green line represents the statistic including all ASVs.

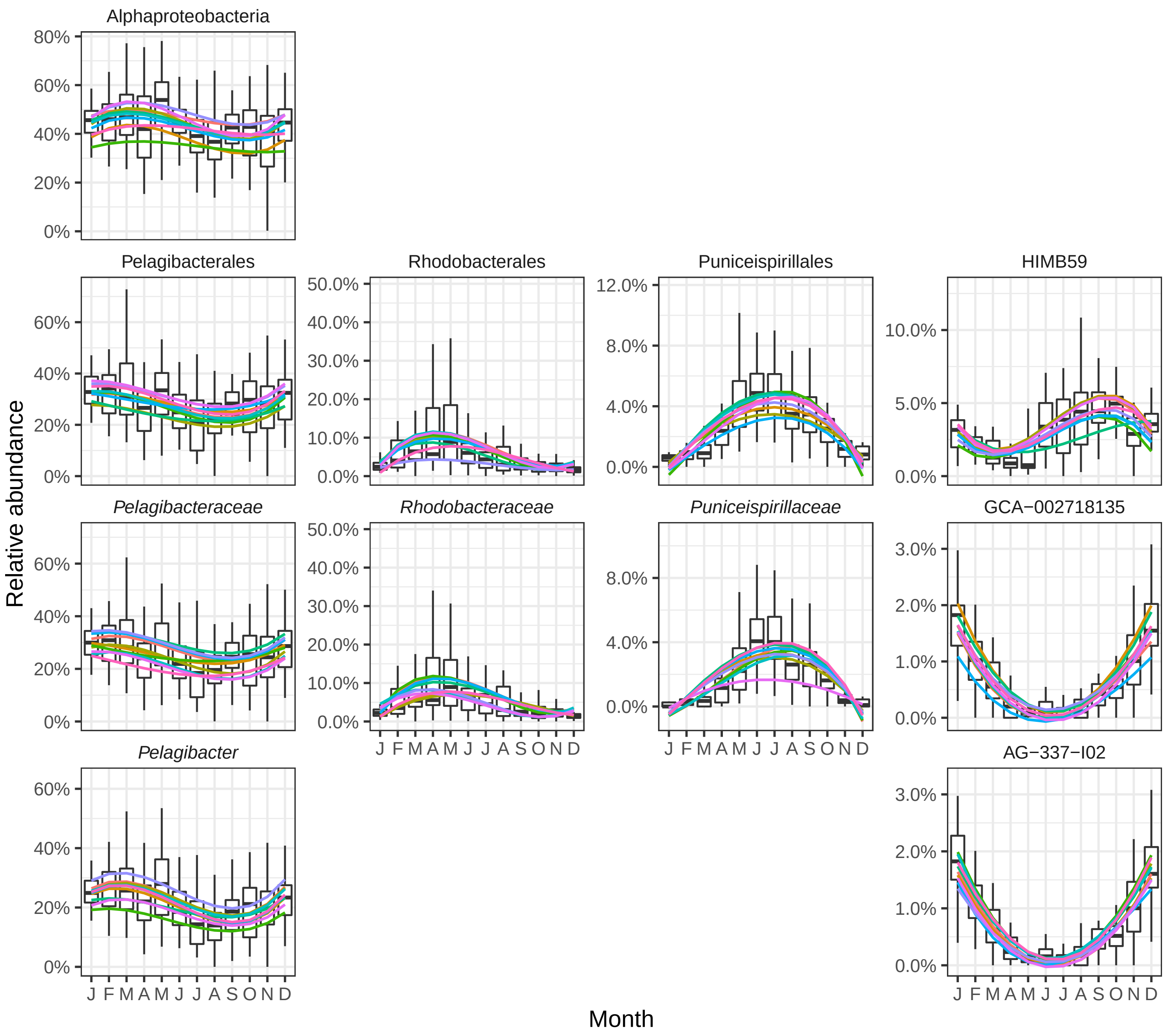

Figure S8: B) Relative abundance distribution of a random selection of 80% of the ASVs calculated 10 times (each line in a different color). Each boxplot presents the median and the 25% and 75% limits of the distribution of 110 points, and whiskers represent 1.5 times the interquartile range. The line is a smooth fitting of the change over time, with a color for each randomization.

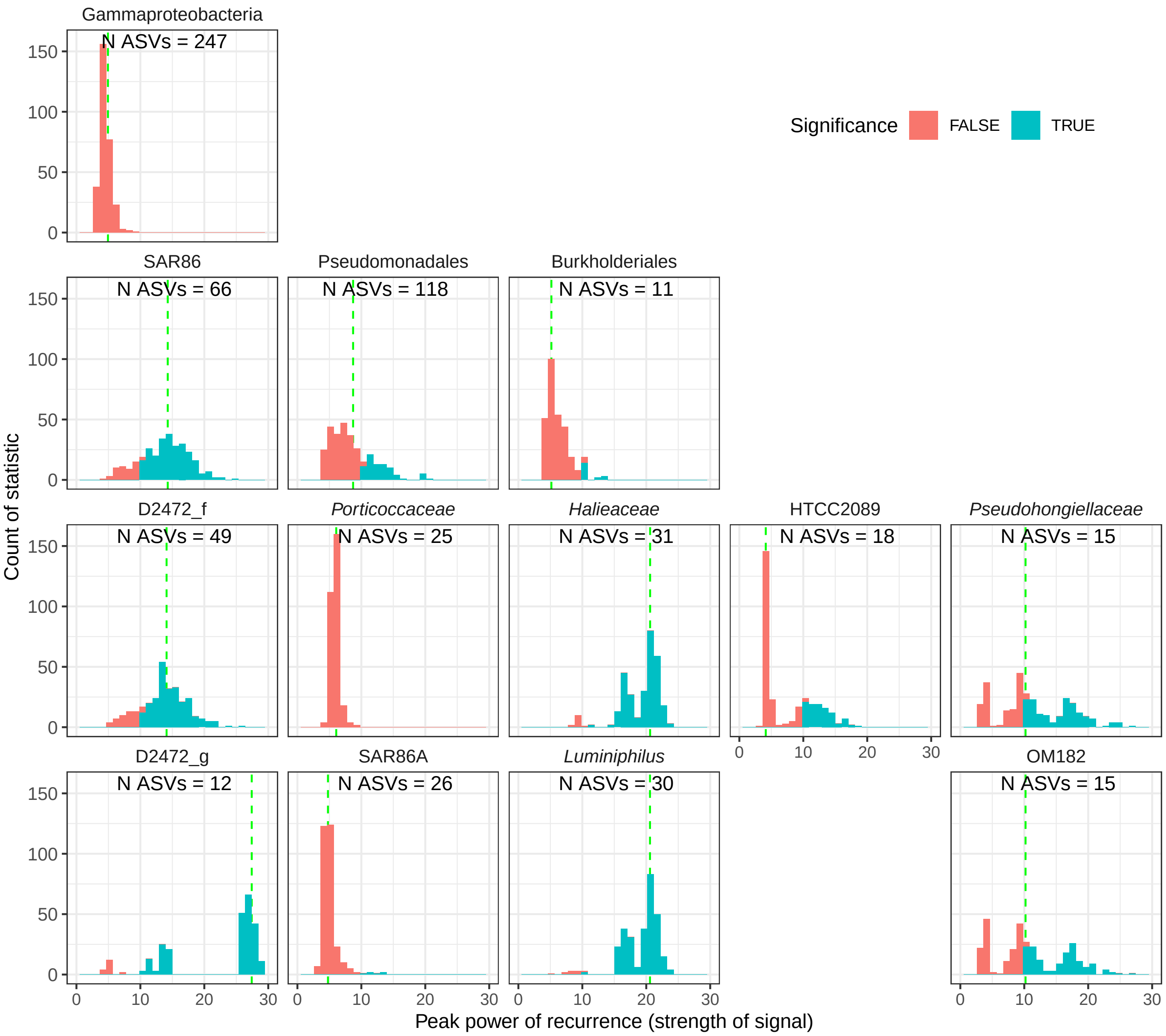

Figure S9: A) Histograms of the peak normalized power statistic at the class, order, family and genus level (from top to bottom, each line is a rank) of 80% of the ASVs conforming the rank for class Gammaproteobacteria. The red bins indicate the non-significant results (PN≥10, $q \leq 0.01$) and blue bins the significant ones. The dashed green line represents the statistic including all ASVs.

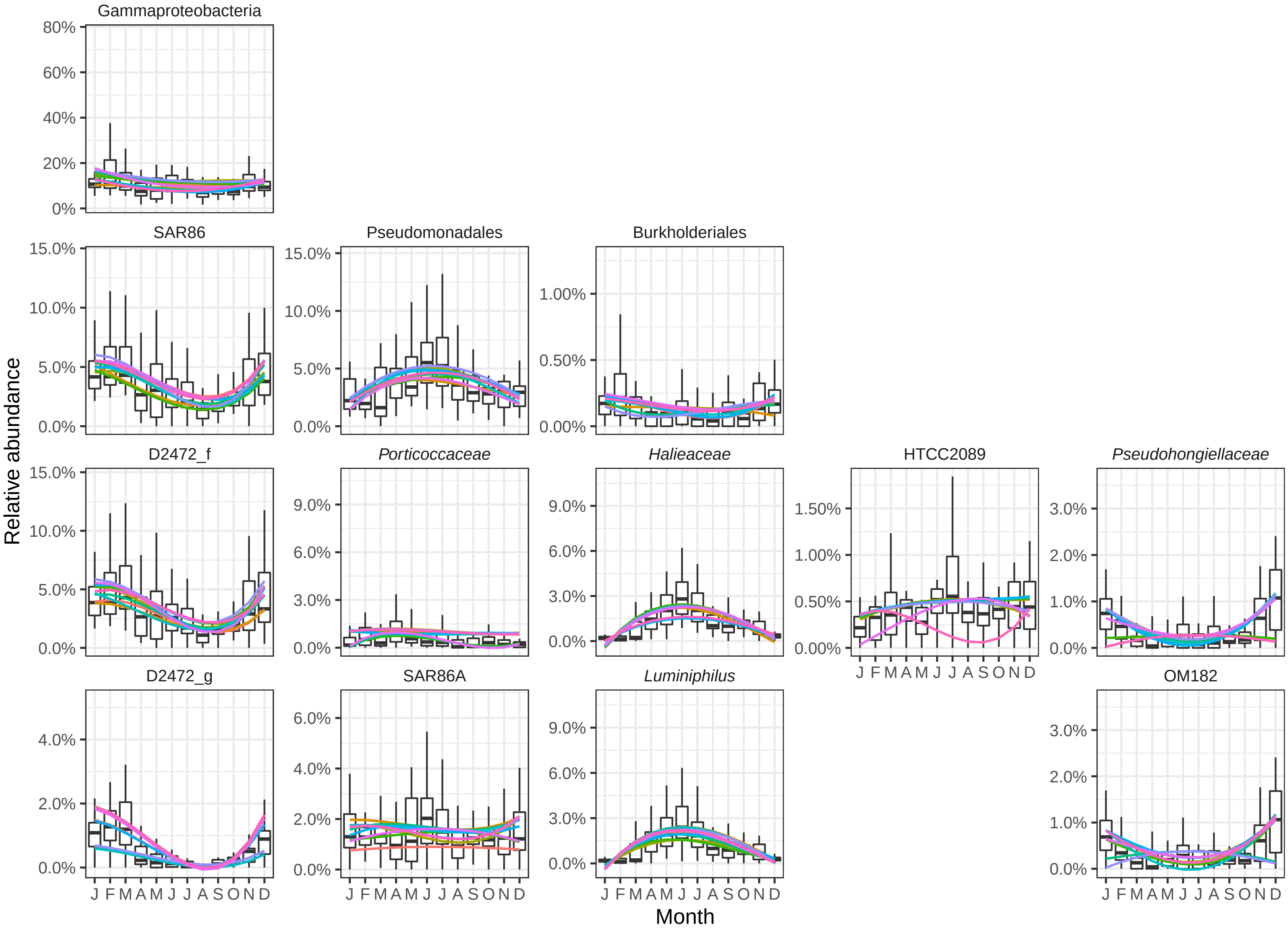

Figure S9: B) Relative abundance distribution of a random selection of 80% of the ASVs calculated 10 times (each line in a different color). Each boxplot presents the median and the 25% and 75% limits of the distribution of 110 points, and whiskers represent 1.5 times the interquartile range. The line is a smooth fitting of the change over time, with a color for each randomization.

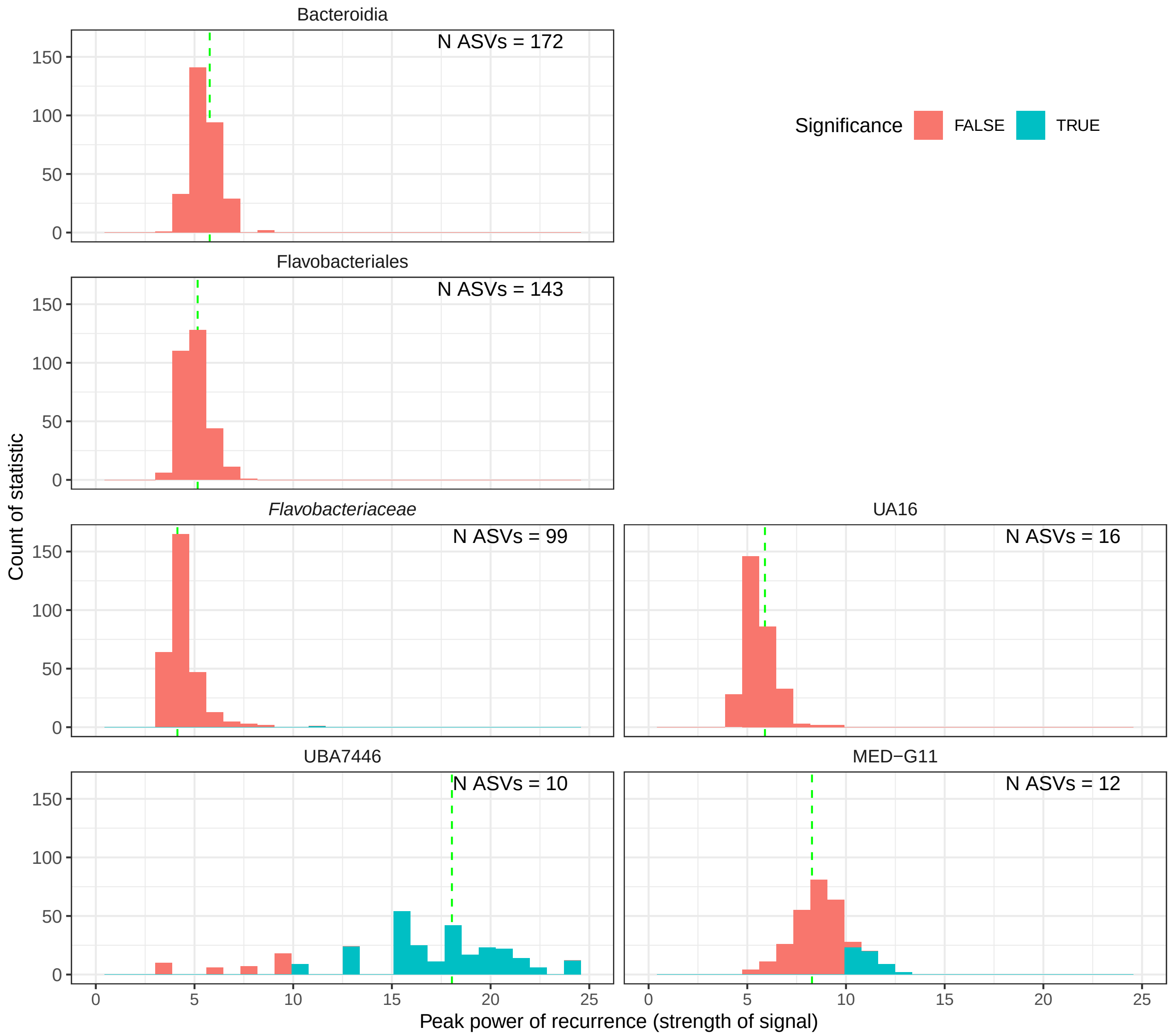

Figure S10: A) Histograms of the peak normalized power statistic at the class, order, family and genus level (from top to bottom, each line is a rank) of 80% of the ASVs conforming the rank for class Bacteroidia. The red bins indicate the non-significant results (PN≥10, $q \leq 0.01$) and blue bins the significant ones. The dashed green line represents the statistic including all ASVs.

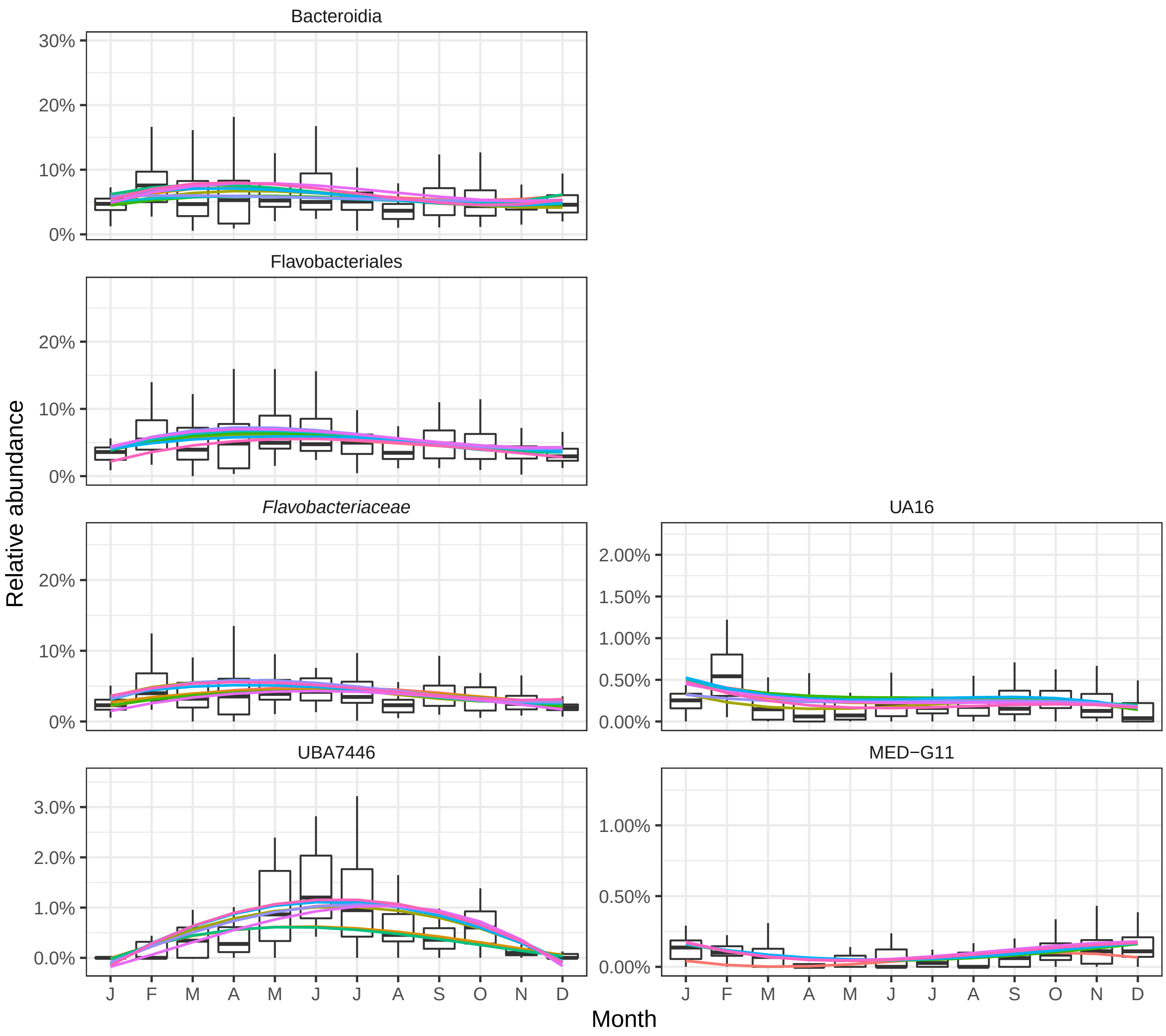

Figure S10: B) Relative abundance distribution of a random selection of 80% of the ASVs calculated 10 times (each line in a different color). Each boxplot presents the median and the 25% and 75% limits of the distribution of 110 points, and whiskers represent 1.5 times the interquartile range. The line is a smooth fitting of the change over time, with a color for each randomization.

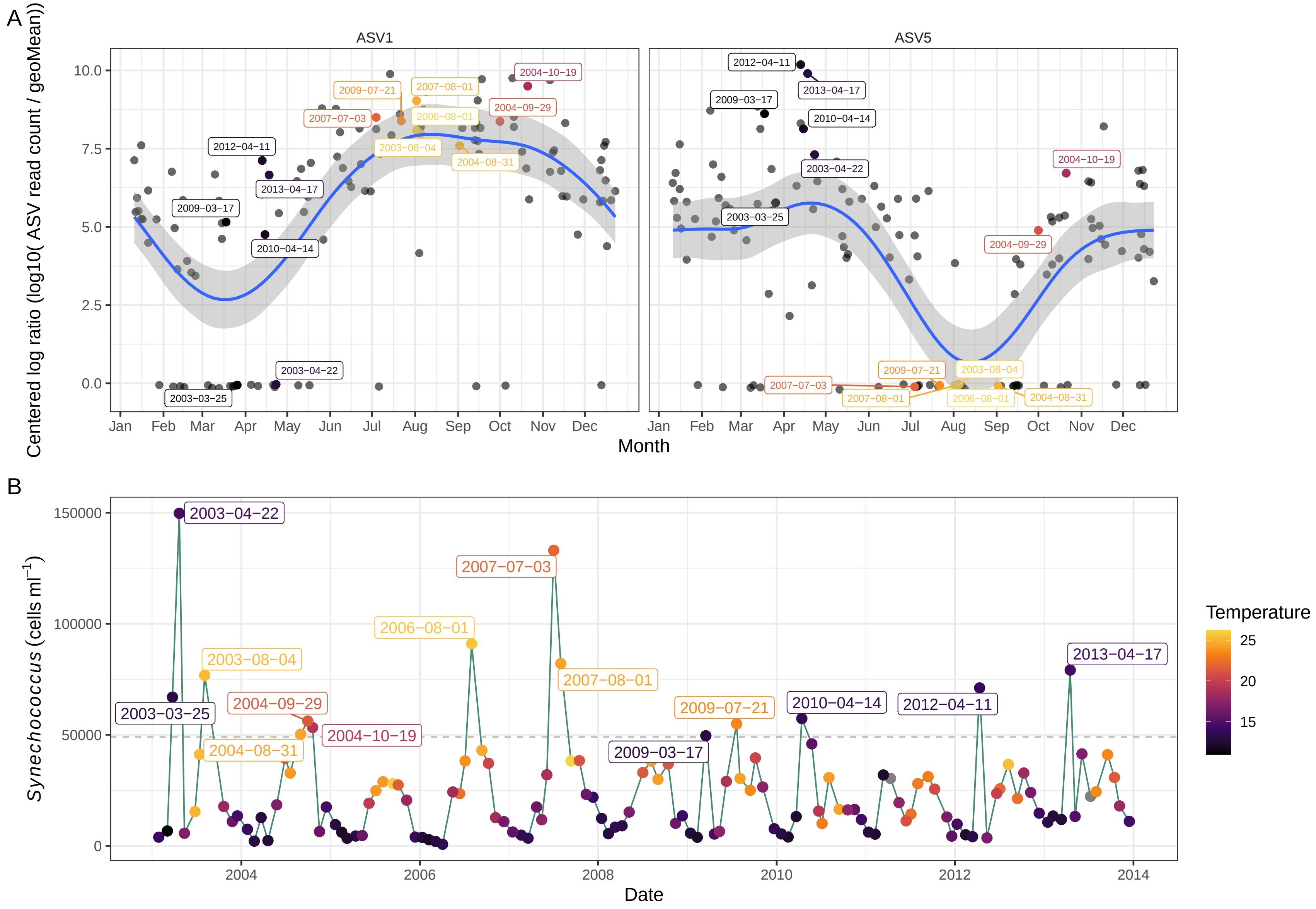

Figure S11: A) Monthly distribution of ASV1 and ASV5, both belonging to the Synechococcus_C genus. The Y axis corresponds to the centered logarithm ratio (with a pseudocount of 1) and the X axis corresponds to the day of the year (month is shown for orientation, with the line ticks for the first day). Dates when a bloom of *Synechococcus* was detected through flow cytometry are labelled. B) Time series of *Synechococcus* abundance (cells $ml^{-1}$) during the 11 years. The data points are colored by water temperature (°C). The grey line indicates the samples presenting more than 50000 cells $ml^{-1}$.
