## Supplementary Table 2 for "Seasonal niche differentiation between evolutionary closely related marine bacteria"

| Genus (GTDB r89) | Order + family (GTDB r89) | Genus (SILVA r138) | Order + family (SILVA r138) | N. seasonal | Total tested ASVs | General information genus |
|---|---|---|---|---|---|---|
| **AG-337-I02** | HIMB59, GCA-002718135 | - | Rhodospirillales, AEGEAN-169 marine group | 12 | 20 | Rhodospirillales order is broken in 4 different orders in GTDB. New studies and data have excluded HIMB59 as a new order outside Rhodospirillales (see *Martijn et al. 2018*). |
| **AG-422-B15** | Pelagibacterales, AG-422-B15 | - | SAR11 clade, Clade IV | 2 | 5 | - |
| **D2472** | SAR86, D2472 | - | - | 3 | 12 | SAR86 order presents 4 families in GTDB: D2472, SAR86, AG-339-G14 and TMED112. In this study we only found assignation for the first family. |
| **HIMB114** | Pelagibacterales, *Pelagibacteraceae* | - | SAR11 clade, Clade III | 4 | 5 | - |
| **HIMB59** | HIMB59, HIMB59 | - | Rhodospirillales, AEGEAN-169 marine group | 2 | 8 | Similar observations to AG-337-I02. |
| **HTCC2207** | Pseudomonadales, Porticoccaceae | SAR92 clade | Cellvibrionales, Porticoccaceae | 0 | 17 | - |
| ***Litoricola*** | Pseudomonadales, *Litoricolaceae* | Litoricola | Oceanospirillales, *Litoricolaceae* | 5 | 8 | - |
| ***Luminiphilus*** | Pseudomonadales, *Halieaceae* | *Luminiphilus* | Cellvibrionales, *Halieaceae* | 9 | 30 | Some assignations included the group in OM60(NOR5) clade for SILVA. |
| ***Marinisoma*** | Marinisomatales, *Marinisomataceae* | - | - | 5 | 8 | Marinisomatota phyla. Not described so far. |
| **MS024-2A** | Flavobacteriales, *Flavobacteriaceae* | NS5 marine group | Flavobacteriales, *Flavobacteriaceae* | 4 | 6 | - |
| **OM182** | Pseudomonadales, *Pseudohongiellaceae* | *Pseudohongiella* | Oceanospirillales, *Pseudohongiellaceae* | 7 | 15 | Oceanospirillales order is included inside Pseudomonadales. |
| ***Pelagibacter*** | Pelagibacterales, *Pelagibacteraceae* | Clade Ia | SAR11 clade, Clade I | 20 | 63 | In some cases the clade was Ib or the assignation was unknown. In this case GTDB unifies instead of splitting as with other groups. |
| **Pelagibacter_A** | Pelagibacterales, *Pelagibacteraceae* | - | SAR11 clade, Clade II | 0 | 27 | - |
| ***Puniceispirillum*** | Puniceispirillales, *Puniceispirillaceae* | Cand. Puniceispirillum | Puniceispirillales, SAR116 clade | 3 | 5 | - |
| **SAR86A** | SAR86, D2472 | - | - | 11 | 26 | - |
| **SCGC-AAA076-P13** | SAR86, D2472 | - | - | 4 | 9 | - |
| **Synechococcus_C** | Synechococcales, *Cyanobiaceae* | *Synechococcus* CC9902 | Synechococcales, *Cyanobiaceae* | 4 | 9 | *Synechococcus* presents 4 genera in GTDB. |
| **TMED189** | TMED189, TMED189 | Cand. *Actinomarina* | Actinomarinales, *Actinomarinaceae* | 4 | 7 | *Actinomarina* assignation is not present in GTDB. Since the assignation comes from *Ghai et al. 2013* this will probably be corrected in further versions of the DB. |
| **UBA4421** | Pseudomonadales, HTCC2089 | - | - | 2 | 7 | - |
| **UBA7446** | Flavobacteriales, *Flavobacteriaceae* | NS4 marine group | Flavobacteriales, *Flavobacteriaceae* | 3 | 10 | - |
