## Supplementary Table 3 for "Seasonal niche differentiation between evolutionary closely related marine bacteria"

| genus | df | logLik | AIC | BIC | deviance | df.residual | pval.term | R.square |
|---|---|---|---|---|---|---|---|---|
| *Pelagibacter* | 2 | 171.5 | –337.1 | –325.2 | 9.1 | 380 | <0.0001 | 0.126 |
| SAR86A | 2 | 15.2 | –24.3 | –21.0 | 0.3 | 20 | 0.052 | 0.135 |
| *Litoricola* | 2 | 14.3 | –22.6 | –19.9 | 0.2 | 16 | 0.683 | –0.051 |
| Pelagibacter_A | 2 | 18.7 | –31.5 | –25.2 | 1.8 | 57 | 0.003 | 0.130 |
| Synechococcus_C | 2 | 2.6 | 0.8 | 2.7 | 0.6 | 12 | 0.89 | –0.082 |
| *Luminiphilus* | 2 | 18.2 | –30.4 | –26.6 | 0.4 | 25 | 0.13 | 0.053 |
| AG-337-I02 | 2 | 11.3 | –16.6 | –13.4 | 0.5 | 20 | 0.19 | 0.038 |
